# Kinetic principles of chemical cross-link formation for protein-protein interactions

**DOI:** 10.1101/2024.02.19.580987

**Authors:** Kai-Michael Kammer, Riccardo Pellarin, Florian Stengel

## Abstract

Proteins play a central role in most biological processes within the cell and deciphering how they interact is key to understand their function. Cross-linking coupled to mass spectrometry is an essential tool for elucidating protein-protein interactions. Despite its importance, we still know surprisingly little about the principles that underlie the process of chemical cross-link formation itself and how it is influenced by different physico-chemical factors. To understand the molecular details of cross-link formation, we have set-up a comprehensive kinetic model and carried out simulations of protein cross-linking on large protein complexes. We dissect the contribution on the cross-link yield of parameters such as amino acid reactivity, cross-linker concentration, and hydrolysis rate. Our model can compute cross-link formation based solely on the structure of a protein complex, thereby enabling realistic predictions for a diverse set of systems. We quantitatively show how cross-links and mono-links are in direct competition and how the hydrolysis rate and abundance of cross-linker and proteins directly influence their relative formation. We show how cross-links and mono-links exist in a “all-against-all” competition due to their simultaneous formation, resulting in a non-intuitive network of inter-dependence. We show that this interdependence is locally confined and mainly limited to direct neighbors or residues in direct vicinity. These results enable us to identify the optimal cross-linker concentration at which the maximal number of cross-links are formed. Taken together, our study establishes a comprehensive kinetic model to quantitatively describe cross-link formation for protein-protein interactions.

**Significance Statement:** Cross-linking mass spectrometry is an essential tool for elucidating protein-protein interactions, but to date basic principles of the process of cross-link formation are still not understood. We set-up a comprehensive kinetic model and simulate the process of cross-link formation. We quantitatively show how cross- and mono-links are in direct competition and how hydrolysis, concentration of cross-linker and of proteins influence their formation. Taken together, our study quantitatively describes kinetic principles of chemical cross-link formation and identifies optimal conditions for cross-link formation.

## Introduction

Proteins assemble into large complexes that play a central role in many biological processes within the cell. Cross-linking coupled to mass spectrometry (XL-MS) is increasingly recognized as an important tool for elucidating protein-protein interactions (PPIs) and for determining the architecture of macromolecular complexes ^1,2^. The general approach of XL-MS is to introduce covalent bonds between proximal functional groups of proteins or protein complexes in their native environment by cross-linking reagents. The actual cross-linking sites are subsequently identified by mass spectrometry (MS) and reflect the spatial proximity of regions and domains within a given protein (intra-link) or between different proteins (inter-link). Additionally, a cross-linker can react only on one side with the peptide and hydrolyze on the other side (mono-link), revealing information on the accessibility of a specific amino acid residue. Over the last couple of years, the field has seen significant technological and conceptual progress and by now the structural probing of recombinantly expressed protein complexes is firmly established. Recent applications of XL-MS on the systems level and in living cells have spurred great interest and hint at the exciting prospect that XL-MS will soon be able to facilitate the structural interrogation of interaction partners of any protein of interest within living cells or even organisms ^3,4^.

As exciting as these recent advancements are and even though an impressive number of increasingly elaborated protein-protein interaction networks are elucidated by XL-MS, we still know surprisingly little about the principles that underlie the process of cross-link formation itself and to what extent different physico-chemical factors contribute to the establishment of a protein cross-link.

Thus, to better understand the factors that contribute to the process of chemical cross-link formation and to quantify their relative contribution we have set-up a comprehensive model of the cross-link reaction based on chemical kinetics and use it to quantitatively simulate the process of cross-link formation. Our kinetic model allows us to define parameters such as the lysine reactivity, the cross-linker concentration, and the hydrolysis rate and enables the detailed investigation of their relative contribution to the overall cross-link yield. The model can also be used to study the cross-link formation based on the structure of any protein complex, thereby enabling realistic predictions of cross-link yields for a diverse set of systems. This is an important point as it often assumed that a distance change, for example by a conformational change that brings two cross-linked residues in closer proximity, will directly result in a change on cross-link abundances, i.e., an increase of the respective cross-link yield ^5–17^. Our model now puts us into a position to directly test this hypothesis. For our model we define a set of parameters which orient themselves at literature values, allowing us to explore the parameter space around these values and to identify a regime under which mono- and cross-link concentrations are maximal, even though the parameter space is in principle of unlimited size. This allows us to explore the parameter space for a “cross-link reaction” in general and to identify conditions that result in maximal cross-link formation and optimal cross-link yields.

For the first time, we show how under standard experimental conditions, cross-links and mono-links within a protein complex are in a “all-against-all” competition due to their simultaneous formation on a protein complex, resulting in a non-intuitive network of inter-dependence. We show that this interdependence is locally confined and mainly limited to direct neighbors or residues in direct vicinity. Our study also describes optimal conditions for cross-link formation from first principles and highlights a way forward in how to design future experiments and new cross-linkers in order to maximize cross-link yields for the study of PPIs. As such it illustrates how to gain access to a larger part of the proteome and helps to pave the way in establishing XL-MS as an essential tool in cellular structural biology.

## Results

### A complete chemical kinetics model of the cross-link reaction

To study the process of cross-link formation and to better understand how different factors contribute to the establishment of a protein-protein cross-link, we have set up a comprehensive kinetic model. We used a homobifunctional, amine-reactive cross-linker (such as NHS-ester based cross-linkers like DSS, DSSO or BS3) as a model system, as the use of chemical groups that react with primary amines are the simplest and by far the most used technique for protein-protein cross-linking. In a first step we described all relevant reactions that occur during the chemical cross-linking of two lysine residues, resulting in a detailed kinetic model which explicitly accounts for the diffusion of all the species (**Fig. 1**; *see* **Materials & Methods** for details). In a second step we then derived an approximated kinetic model, which contains less parameters and is more efficient. A kinetic model is a set of time differential equations which allows for a complete macroscopic description of reactions without the prerequisite of detailed knowledge of all the involved chemical species ^18^. Here, the chemical rate constants represent the frequency of pairwise collisions of reactants that lead to the formation of products, thus our rate equations are first- or second order with respect to the concentration of the species. Integration of our kinetic model over time enables us to simulate the concentration change of all species in a time-resolved manner. Importantly, the parameters of the model (i.e., the rate constants) are all, in principle, experimentally available physical properties.

**Figure 1.**
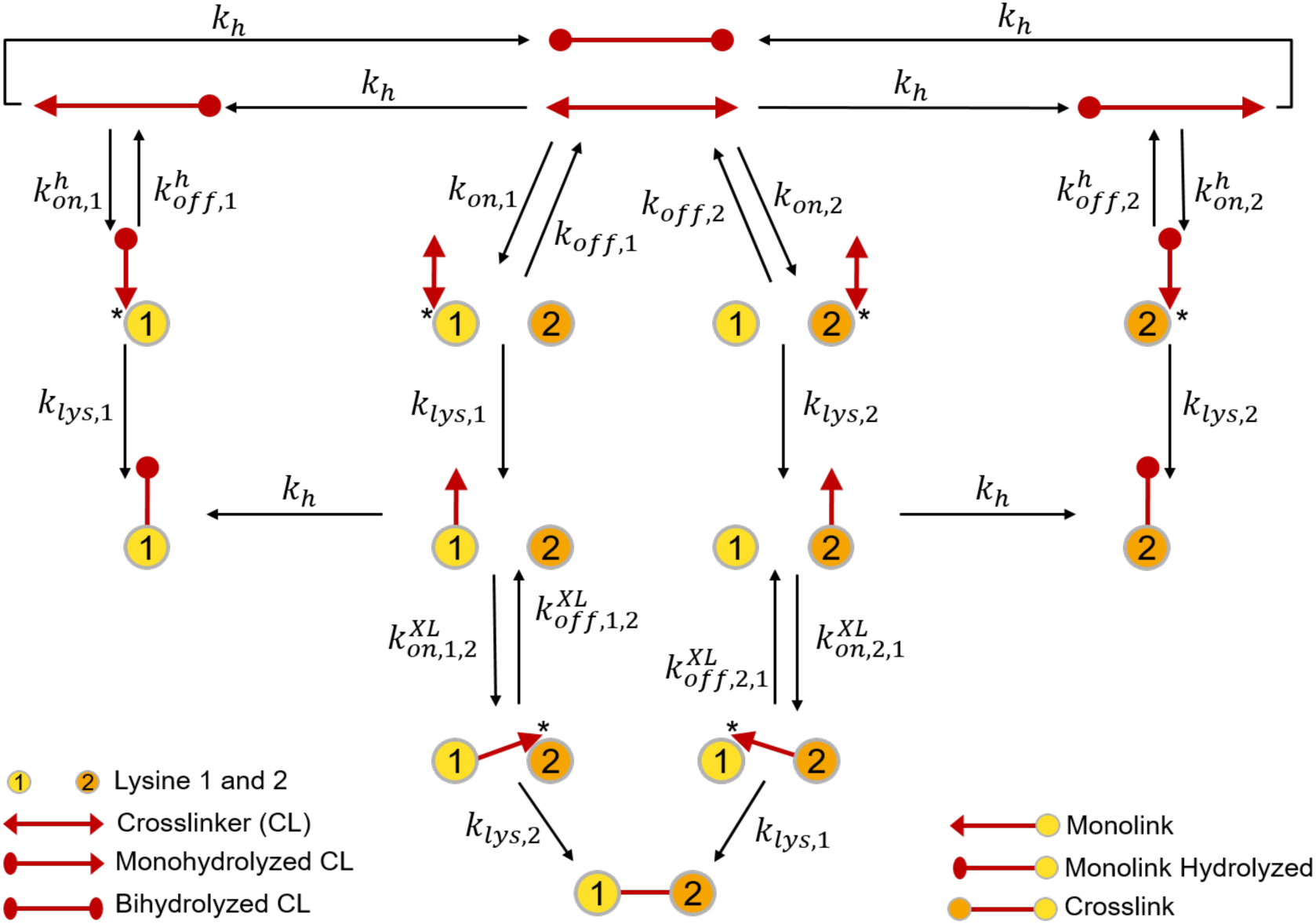
Overview of the full kinetic model of the cross-link reaction. The kinetic model of the cross-link reaction includes hydrolysis, mono-link formation, and cross-link formation. A cross-linker may hydrolyze twice: first forming a mono- and then a bihydrolyzed cross-linker. The monohydrolyzed cross-linker may still form a mono-link, while the bihydrolyzed cross-linker is a dead-end of the reaction. To form a cross-link, a non-hydrolyzed cross-linker needs to first form a mono-link. This mono-link then needs to encounter another lysine in proximity to form a cross-link. Shown is a two-lysine model as an example, even though the model is generalizable to any number of reactive sites. The model is fully described by 15 unique rate constants which are depicted above the reaction arrows. *k_on_* and *k_eff_* describe diffusion rates, *k_h_* hydrolysis and *k_lys_* lysine reactivity. Of special importance is *k^XL^_on_* which represents the distance between a lysine pair as a diffusion rate. We call this rate the *kinetic distance* (*see* **Materials & Methods** for details).

Our model accounts for three main reactions: 1) hydrolysis, 2) mono-link formation, and 3) cross-link formation. Hydrolysis of reactive groups of the crosslinker plays an important role. It leads to “dead-ends” in the reaction pathway, resulting in species that cannot react any further to form crosslinks, i.e., the reaction stops in a bihydrolyzed cross-linker or a hydrolyzed mono-link. Hydrolysis depletes free cross-linker and, as a consequence, decreases the amount of formed mono- and cross-links. Mono-link formation can occur via two different pathways and starts either with a nonhydrolyzed or a monohydrolyzed cross-linker. Likewise, a cross-link can also arise from two parallel pathways, where the two possible precursor mono-links act as intermediates (**Fig. 1**). In our model all reactions are described by rate constants. We directly relate the structure of a system, i.e. a protein complex, with a rate constant by encoding the distance between two cross-linked residues into a diffusion rate constant of the cross-link reaction (termed “*kinetic distance”)*. We also define other physico-chemical parameters like the intrinsic lysine reactivity, cross-linker amount, diffusion, and hydrolysis rate. All these parameters allow us to *quantitatively* dissect their relative contribution to the overall cross-link yield. From this detailed model we then derive an approximated kinetic model, which consists of less parameters and is more efficient. This simplified kinetic model (**Suppl. Fig. 1**) aggregates the diffusion of the species into effective rates, thereby reducing the number of free parameters. All the following simulation results are based on this simplified model. As our model is scalable to any number of lysines, we developed a python framework that uses solely the structure (pdb file) as input to generate the parameters and the equations of the kinetic model for a given protein. In a final step a model is created for each protein and parameter combination, which can then be simulated using a kinetic simulation backend (*see* **Materials & Methods** for details). Taken together, we have established a comprehensive kinetic model that allows us to quantitatively describe the process of cross-link formation.

### Mono- and cross-links are in direct competition

Simulations of our kinetic model revealed the existence of three separate forms of competition between the different species that are formed during a crosslinking reaction: 1) competition between mono-links and cross-links, 2) competition between different cross-links and 3) competition between mono-links and cross-links on one side and hydrolysis on the other.

To better understand the principles of this competition, we have set-up a simple kinetic model consisting of only three lysine residues, labeled A, B, and C, that are placed at the same kinetic distance, such that each mono-link has the same probability to collide with a free lysine to form a cross-link (**Fig. 2**; *see* **Materials & Methods** for details). We then carried out three sets of simulations. In the first, the reactivity of all lysine residues was assumed to be identical, while in the other two we modulated the reactivity of the lysines. We used the final cross-link and mono-link concentrations of the first simulation as reference to quantify relative changes due to perturbations of the system introduced in the other two simulations (**Fig. 2A**, *upper row*). As expected, all three mono- and cross-links had the same concentration in this first simulation. In the second simulation, the reactivity of lysine A was decreased by one order of magnitude. Here, we observed a decrease in the concentration of the corresponding mono-link as well as a decrease in the concentration of the two directly involved cross-links (A-B & A-C). Interestingly, we also find an enrichment of the third cross-link between the two lysine residues with unchanged reactivity (B-C) as well as increased mono-link concentrations on lysine residues B and C (**Fig. 2A**, *middle row*). In the third simulation, the reactivities of two lysines (A and B) were decreased by one order of magnitude. Here, we observe that the corresponding mono-links (A and B) were depleted, as well as all cross-links, with the largest reduction taking place at the cross-link between lysine residues A-B. Interestingly, mono-link concentrations for lysine C, whose reactivity was not changed, was enriched by a factor of 1.5 with respect to the first simulation (**Fig. 2A**, *bottom row*).

**Figure 2.**
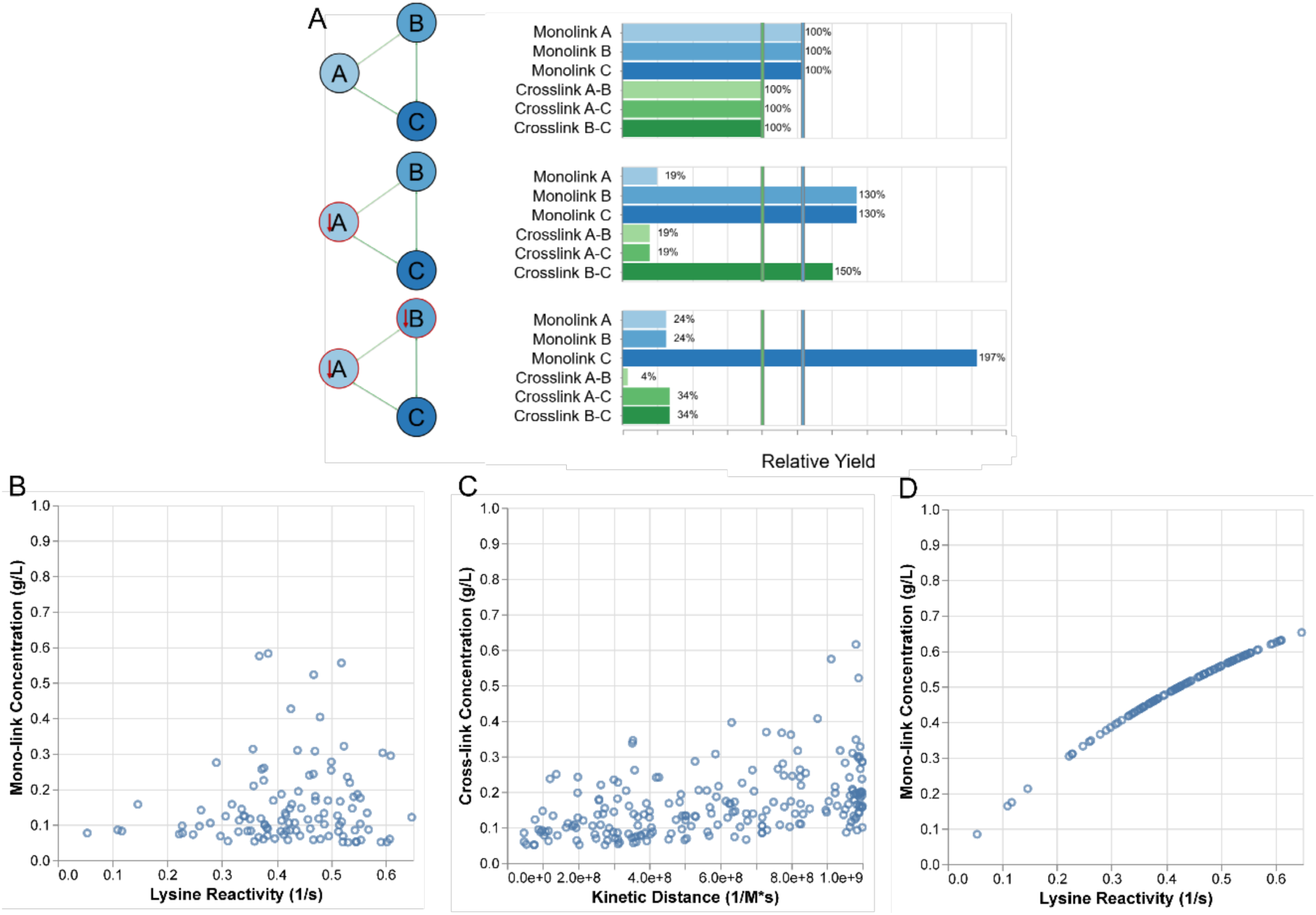
Mono-links and cross-links are inter-dependent. **A** Results of a three-lysine simulation. *Top:* results for all lysines having the same reactivity. *Middle:* the reactivity of lysine A is reduced by a factor of ten. *Bottom:* both lysine A and B have their reactivity reduced by a factor of 10. The cross-link concentrations are shown in green and the mono-link concentrations in blue. The first row is used as reference as indicated by the blue and green bar for mono- and cross-links, respectively**. B** and **C** are the results obtained for a standard simulation, where cross-links are allowed to form together with mono-links on a structure of the C3 complement factor. Each dot represents the final concentration of a mono- or a cross-link. **B** The final concentration of mono-links is plotted against the lysine reactivity. **C** The final concentration of cross-links is plotted against the kinetic distance (*see* Materials and Methods for details). **D** The final concentration of mono-links is plotted against the lysine reactivity for an ideal “mono-link-only” simulation, i.e. where no cross-link reaction can take place, using the structure of the C3 complement factor. *See* **Supp** Fig 2 for an overview including all studied model protein complexes.

Taken together, this simple set of simulations clearly demonstrates that concentrations of neighboring mono- and cross-links are co-dependent, if they share the same lysine. A change in reaction rate for one lysine residue will therefore propagate and affect the reaction rate for all mono- and cross-links competing for that specific lysine.

### Mono- and cross-link abundances are inter-dependent

The observed competition between cross-links and mono-links in our simple three-lysine simulation suggests, that such an interdependence of mono- and cross-link yields should also exist on a larger scale. However, such an interdependence would imply that the *distance* between two cross-linked lysine residues *cannot* be the sole factor that determines relative crosslink yields, as a change in lysine reactivity and mono-link concentration could affect cross-link quantities, directly or indirectly. This is an important point as it is often assumed that a distance change, for example by a conformational change that brings two cross-linked residues in closer contact, will directly result in a change of cross-link abundances, i.e., an increase of the respective cross-link ^5–17^. Our model enables us to directly test this hypothesis, as one would expect a strong correlation between the final quantity (i.e. concentration) of a formed cross-link and the kinetic distance of the corresponding lysines, if a cross-link reaction was indeed mainly controlled by the distance between the linked lysine-residues. We therefore set up a model system based on the six proteins and protein complexes Eg5, Luciferase (Luci), the complement factor C3, RNA polymerase II (RNAP), Human IFT-A complex (IFTA) and the 26S proteasome (**Table 1**). These proteins and protein complexes were selected because they cover a wide range in terms of molecular weight and exhibit a large variety in terms of their 3D structure and lysine distribution.

**Table 1.** Proteins complexes and their PDB IDs used for the kinetic simulations, with their corresponding simulated concentrations.

| Protein | PDB ID | Molecular Weight<br>/kDa | No. Lysines | Concentration (at<br>1 g/L)<br>mol/L |
| --- | --- | --- | --- | --- |
| Eg5 | 3WPN | 43 | 21 | 2.31e-5 |
| Luciferase | 4G36 | 122 | 75 | 8.21e-6 |
| C3 | 2A73 | 184 | 111 | 5.41e-6 |
| RNAP | 6DRD | 523 | 203 | 1.91e-6 |
| IFTA | 8FGW | 767 | 381 | 1.31e-6 |
| 26S | 4CR2 | 1308 | 749 | 7.71e-7 |

However, when subjecting these protein model systems to our cross-link simulations, we find no direct correlation between the kinetic and the physical distance (**Fig. 2B & 2C; Supp Fig 2**). We see also no discernible correlation between the reactivity of a lysine and the corresponding mono-link quantity. While this does not contradict the notion that a distance change may lead to a change in cross-linking quantity, it clearly demonstrates that distance is not the sole determining factor for cross-link quantity.

This picture changes dramatically when we are looking at a “mono-link only” situation, i.e. a simulation where the reaction stops with the mono-link reaction and is not allowed to proceed along the cross-link reaction pathway. In this simulation, we see a strong correlation between mono-link quantity and corresponding lysine reactivity (**Fig. 2D**; **Supp Fig 2**). We therefore conclude from this data, that it is indeed due to the competition between cross-link and mono-link reaction that this correlation is lost.

To study this important observation further, we systematically analyzed how mono-link and cross-link quantities are influenced when the reactivity of a given lysine is suppressed. We therefore consecutively suppressed (i.e. set to 0) the reactivity of each lysine in one of our model proteins (**Supp Fig 3A**). In a next step we then monitored the number and strength of significant changes in mono- and cross-link quantity that were caused by this suppression (**Supp Fig 3B**). We find that significant changes are mainly limited to residues that are located very closely to each other (**Supp Fig 3C**). For example, for the Eg5 protein model we find that suppression of lysine at position 146 influences the yield of three cross-links and two mono-links (**Supp Fig 3D**), which were all directly connected via a cross-link to lysine 146 before the suppression.

Taken together, our data shows that a simple correlation between physical distance and abundance of cross-linked residues does not exist due to the complex competition between cross-links and mono-links. However, our data also demonstrates that this interdependence is locally confined and mainly limited to direct neighbors or residues in direct vicinity.

### Mono-and cross-link concentrations are affected by hydrolysis in different ways

Hydrolysis is in constant competition with the mono-as well as the cross-link reaction for free cross-linker. To investigate how hydrolysis quantitatively affects mono-link and cross-link formation we ran cross-linking simulations on our six model protein complexes.

Our model shows that mono- and cross-link concentrations are differently affected by a change of hydrolysis strength (**Fig. 3**). We find that a peak in the concentration of mono-links corresponds to a decrease in cross-link concentration. This effect can be explained by the fact that cross-links are more affected by hydrolysis than mono-links because the cross-link reaction occurs in two steps, where the mono-link is a compulsory intermediate. The cross-link reaction is a two-step 2^nd^ order reaction with respect to both the concentration of a given lysine and a given mono-link. The mono-link reaction on the other hand can be described in much simpler terms as a one-step 2^nd^ order reaction, which solely depends on the concentration of a specific lysine residue and the overall cross-linker concentration (for details *see* **Materials & Methods**). Thus, there are two possible ways for hydrolysis to stop the cross-link reaction and to prevent the formation of a cross-link.

**Figure 3.**
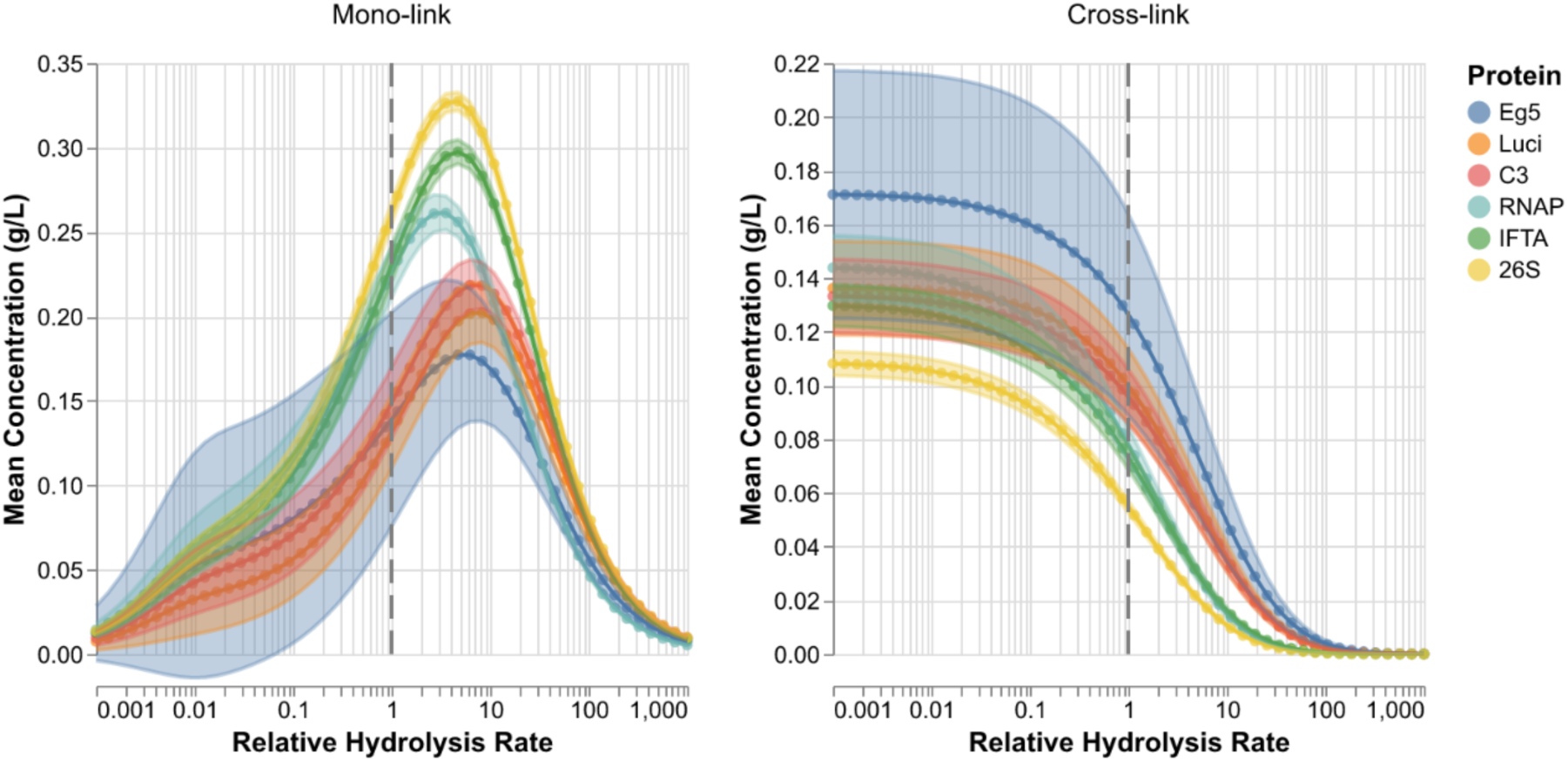
Mono-and cross-link concentrations are differently affected by hydrolysis. The mean mono-link (solid circled lines, left) and mean cross-link (solid circled lines, right) concentration plotted against the relative hydrolysis rate for six model proteins (sorted from smallest to largest by molecular weight). The reference hydrolysis rate used throughout this study is 2.5e-4 s-1 and is indicated by the dashed line (**Supp Table 1**). All other parameters are kept constant. The colored band represents the variance of the final concentration computed over all different species.

This results in the non-trivial effect that *increasing hydrolysis* leads at first to a *higher mono-link concentration* (**Fig. 3**, *left panel*). As a necessary intermediate, mono-links “profit” from cross-links that are more affected by an increased hydrolysis rate, resulting in a maximum in mono-link concentration at a medium-hydrolysis rate. As expected, if the hydrolysis rate is increased even further, this eventually leads to a decrease of both species.

Hydrolysis has another remarkable effect on how protein concentration affects the ratio between mono- and cross-links. Both mono- and cross-links depend on the concentration of available lysine residues and a lower molar lysine concentration means slower rates for both mono- and cross-link formation. In principle, both reactions should be slowed down equally when lysine and cross-linker concentrations are reduced proportionally. However, as described above, hydrolysis affects cross-links more than mono-links and this effect becomes particularly apparent at low protein concentrations (**Supp.** Fig. 4). This observation is for example important in experiments where cross-linking of recombinant proteins and protein complexes is carried out, where usually a fixed mass concentration of typically 1 g/L is used ^19^. This results in a decrease of the molar concentration of a protein or protein complex with increasing size and shifts the ratio between mono- and cross-links in favor of the former. We can also observe, that at a fixed molar concentration hydrolysis plays less of role for larger proteins, as hydrolysis is no longer able to compete with the increased amount of lysines and cross-linker (**Supp Fig 4**). On the other hand, this also means that *without* hydrolysis, one could achieve equal mono-to cross-link ratios independent of protein size, given sufficient reaction time (**Supp Fig 4**).

Taken together, we demonstrate how mono- and cross-link concentrations are differently affected by hydrolysis. We show that under conditions of lower protein concentrations mono-links will be favored over cross-links and how hydrolysis is a tunable parameter that can be used to shift the overall reaction towards mono- or cross-links.

### Mono-and cross-link concentrations are differently affected by cross-linker concentration

In the next step we used our model to test how the cross-linker concentration quantitatively affects mono-link and cross-link formation on our set of model protein complexes, as it is one of the experimentally most readily accessible parameters.

As expected, we find that the mono-link concentration continuously increases with increasing cross-linker concentration (**Fig 4**, *left panel*). However, to our surprise, we find that the cross-links did not follow the same trend. We observe that the cross-link concentration follows a similar trend as the mono-link concentration, but only up to a certain maximum level, depending on the model parameters (**Fig 4**, *right panel*). This peak point is reached for all our protein models and in all cases corresponds to a half-equimolar to equimolar cross-linker-to-lysine concentration. Beyond this maximum the cross-link concentration starts to decrease again.

**Figure 4.**
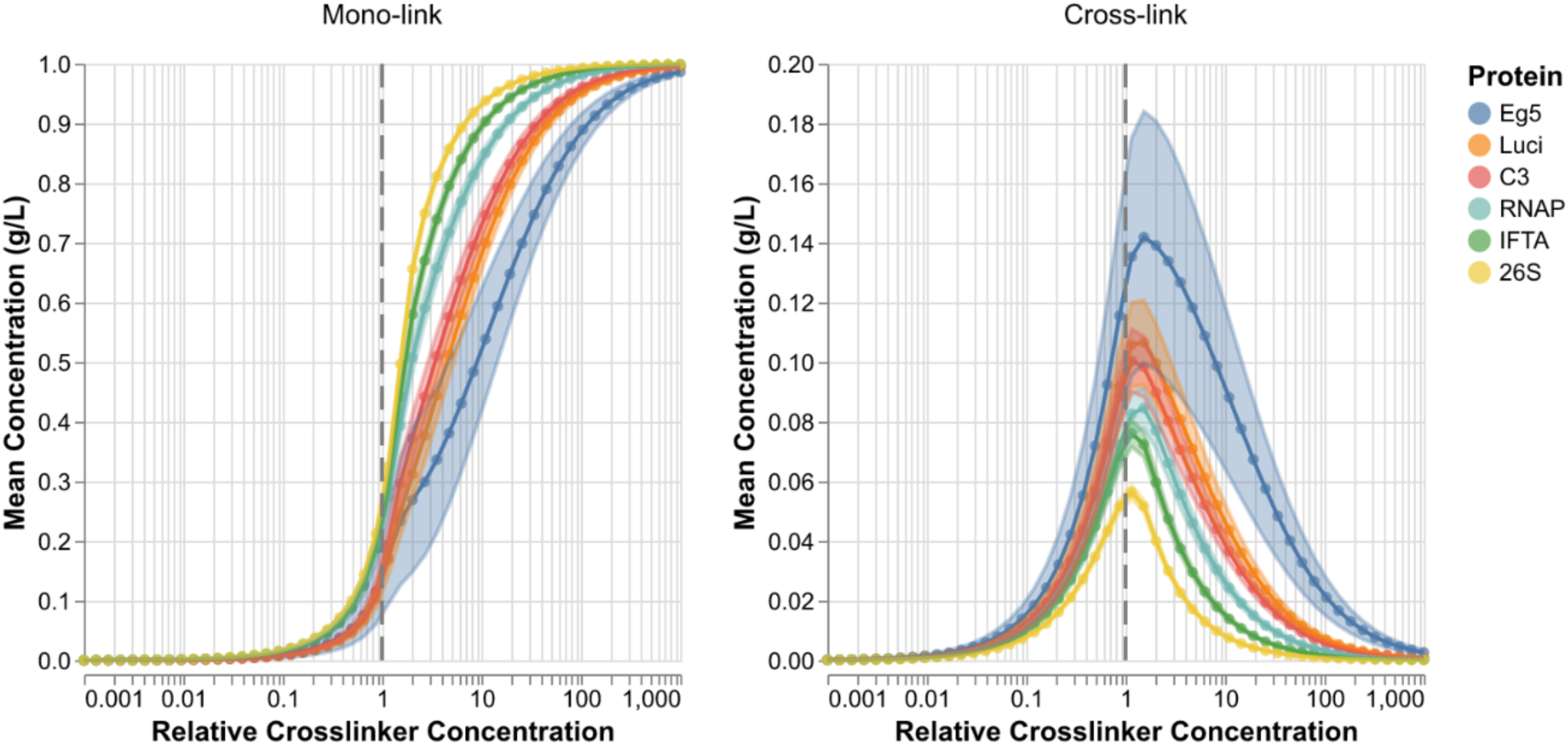
Mono-and cross-links are differentially affected by the cross-linker concentration. Mean mono-link (solid circled lines, left) and mean cross-link (solid circled lines, right) concentration plotted versus the relative cross-linker concentration for six model proteins (sorted from smallest to largest by molecular weight). The reference cross-linker concentration used throughout this study is 0.5 times the number of lysines in g/L and is indicated by the dashed line (**Supp Table 1**). All other parameters are kept constant. The colored area represents the variance of the concentration computed over all species.

This at first sight surprising result is readily explained if we examine the cross-link reaction on a microscopic scale. If two lysine residues that could in principle form a cross-link have already reacted with a cross-linker, i.e. have both already formed a mono-link, they can no longer react in a second step to form a cross-link. The higher the cross-linker concentration, the higher is therefore the probability that two given lysine residues in proximity are both already mono-linked *before* they have the chance to react further to form a cross-link. Under such high cross-linker concentration conditions, mono-links are therefore favored over cross-links.

Taken together we therefore find that there exists an optimal cross-linker concentration at which the maximal number of cross-links can be formed and that this optimal cross-linker concentration is at a half-equimolar to equimolar cross-linker-to-lysine concentration.

### Multi-parameter influence on cross-link formation

Next, we wanted to study how variations in a multiple parameter space influence cross-link formation. Therefore, we explored the effect of varying cross-linker and lysine concentrations simultaneously, as well as varying cross-linker concentration and hydrolysis strength, to then show the amount of mono- and cross-cross-links that have formed at the end of the reaction in a phase diagram (**Fig 5**). We observe that if we fix the lysine concentration (which we can determine from the protein concentration and the number of lysines in our respective model protein) and if we then increase the cross-linker concentration, mono-links start to become even more favored over crosslinks (**Fig 5A and Fig 5B**). Here, it is also interesting to note that due to hydrolysis we do not see any significant number of formed mono-links in the low lysine or cross-linker concentration regime.

**Figure 5.**
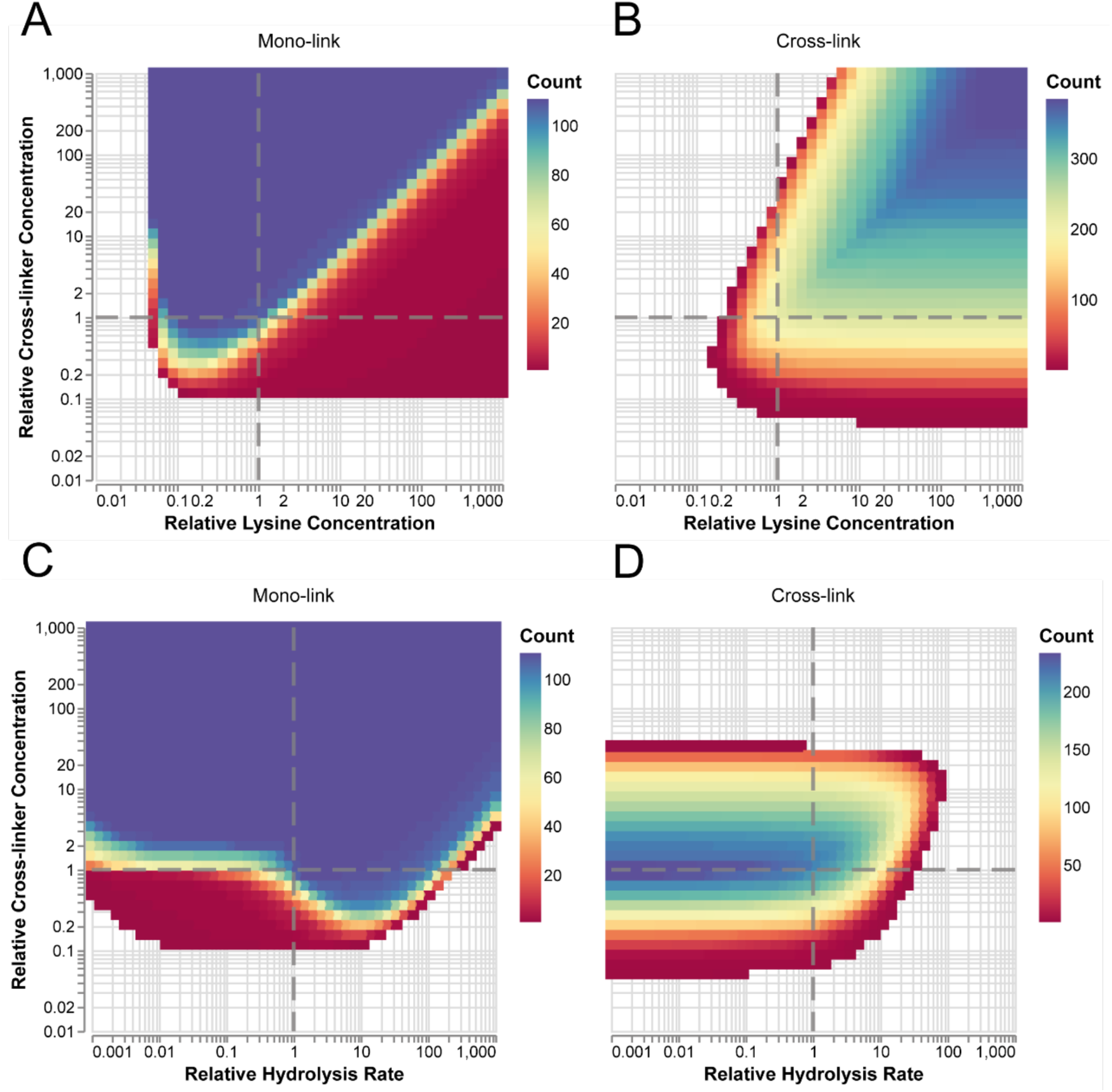
Cross-link and mono-link formation as a multi-parameter function. First row: relative cross-linker concentration plotted versus the relative mean lysine concentration. Second row: relative cross-linker concentration plotted versus the relative hydrolysis rate. The color corresponds to the number of formed mono-links (**A**, **C**) and cross-links (**B**, **D**) identified for the given parameter combination using an intensity cut-off of 0.05 g/L. The reference value of each parameter is indicated by the dashed lines (**Supp Table 1**). Simulation results are shown for the structure of the human complement factor C3. Simulations for additional model proteins are shown in the Supplementary Information (**Supp** Fig 5 and **Supp** Fig 6).

If we shift our focus to cross-links, we observe that the region where the highest number of cross-links are formed (bluish area of the phase diagram) runs along a 45-degree line from the origin (**Fig 5B**). Correspondingly, we find a region of mono-link depletion along the same 45-degree line (as indicated by the transition between the blue and red areas in the phase diagram in **Fig 5A**). This means that a fixed ratio between cross-linker and lysine concentration is required to obtain a maximum cross-link yield, independent of absolute cross-linker/lysine concentrations. As stated above, for low cross-linker/lysine concentrations hydrolysis starts to dominate and prohibits any significant cross-link or mono-link formation (**Fig 5A and Fig 5B**).

If we now vary the cross-linker concentration and the hydrolysis rate simultaneously (**Fig 5C and D**), the number of mono-links is affected only under a regime where mono-link formation is in competition with cross-link formation, i.e. under a regime of lower cross-linker concentration, as can be seen in the lower part of the phase diagram (**Fig 5C**). Under these conditions the region where the highest number of cross-links are formed (bluish area of the phase diagram) is found at a constant cross-linker concentration, i.e. runs along the X-axis (**Fig 5D**), irrespective of the hydrolysis rate. Only for very high hydrolysis rates, both mono- and cross-links are depleted (**Fig 5C and D**).

In summary, we find that high mono-link yields are more readily attainable than high cross-link yields and that mono-links are less affected by hydrolysis. In order to obtain maximal cross-links, one therefore needs to ensure that the cross-linker-to-lysine ratio is at its optimum, i.e. at a half-equimolar to equimolar.

## Discussion

In this work we report on the development of a model and simulation package of the cross-linking reaction based on chemical kinetics. Our model, to the best of our knowledge, is the first complete kinetic description of the cross-link reaction itself. Even though we have tailored it to amine-reactive cross-linkers as a model system, due to their widespread use, the model is easily adaptable to other chemistries. All the parameters of the model (i.e. rate constants) are approximations, but are in principle experimentally available physical properties, and should therefore duly describe the physical world. For our model we could partly build on prior work, where we could demonstrate that kinetics influence the propensity of cross-links to preferentially form on high abundant proteins ^20^, but it is only now that we are in a position to define parameters like lysine reactivity, cross-linker amount and diffusion, and hydrolysis rate in order to dissect their relative contribution to the overall cross-link yield quantitatively.

It is important to note that our model and the corresponding simulations represent *only* the chemical reaction of the cross-link formation. While concentrations are directly obtainable *in silico*, experimental cross-linking data must be obtained by experimental means and depends on the methods and mass spectrometer used. Consequently, single data points can either not be recorded or can be disregarded in the statistical evaluation of a given experiment^21–31^. Therefore, the results presented here showcase the *ideal case* of obtaining data without experimental error.

We show that there is competition between the various species involved in a cross-linking reaction and, with the help of our comprehensive model, we quantitatively assess the relative contribution on each level. Here, our model allows us to clearly dissect how cross-links and mono-links compete amongst themselves, with each other and against hydrolysis for reactive sites (i.e. lysine residues). This results, for example, in the non-trivial effect that increasing hydrolysis strength leads at first to a higher mono-link concentration.

Our model clearly shows that structurally neighboring mono-and cross-links quantitatively affect each other and are interdependent. This observation has important implications, as our model demonstrates that this effect is intrinsically entangled with the effect that links cross-link distance with lysine reactivity. Ultimately, we learn from our model that the reaction rates of all chemical species are interdependent if they are competing for the same lysine. A change in reaction rate for one species will therefore affect the concentrations of all mono- and cross-links competing for that specific lysine, and this propagates to a smaller extent to second neighbors, in a network-like fashion. This, on the other hand, necessitates that distance is not the sole determiner for cross-link quantity; a notion that up to now is widely accepted in field, as it is assumed that a distance change, for example by a conformational change that brings two cross-linked residues in closer contact, will directly result in a change on cross-link abundances, i.e., an increase of the respective cross-link ^5–17^. The simulations of our kinetic model show that if the cross-link reaction was indeed mainly controlled by the physical distance between two linkable residues, one would expect a strong correlation between the final concentrations in our simulation and the kinetic distance. However, we only find a marginal correlation between these two variables and at the same time we see no discernible correlation between the reactivity of a lysine and the corresponding mono-link quantity. This picture changes only once we start to look at a mono-link only simulation, where the reaction stops after the formation of a mono-link. Only in this regime do we see a direct correlation between mono-link quantity and lysine reactivity. It is therefore due to the complexity of the cross-link reaction itself that this correlation is lost. One way forward to probe lysine reactivities and to deconvolute the complex relationships within networks formed by cross-links even further could therefore be the use of *monofunctional* NHS ester, as they would allow us to also test a mono-link only regime experimentally.

Our kinetic model also demonstrates how mono- and cross-link concentrations are differently affected by hydrolysis and how hydrolysis is a tunable parameter that can be used to shift the overall reaction towards mono- or cross-links. It shows that there is a protein concentration-dependence not only for absolute numbers of cross-links and mono-links but also for the cross-link to mono-link ratio, strongly favoring mono-links at lower molar concentrations. This effect is facilitated by hydrolysis and not observable in simulations *without* hydrolysis, where one could achieve equal mono- to cross-link ratios also for lower protein concentrations given sufficient reaction time. This effect is not only relevant for experiments where recombinant proteins or protein complexes are employed and where usually a fixed mass concentration is used. Our data would also predict that the number of cross-links that can be obtained in proteome-wide settings using cross-linking is generally limited, in particular when hydrolysable cross-linkers as amine-reactive linkers are used. However, it also suggests that one way forward to increase the number of PPIs that can be formed by cross-links and to gain access to a larger part of the proteome could be the design and use of non-hydrolysable cross-linkers or the use of UV-activatable caged linkers.

As our comprehensive kinetic model allows us to explore not only any parameter in detail but multiple parameters in parallel, we are in the unique position to universally explore the parameter space for a “cross-link reaction”. Here, we find that there exists an optimal cross-linker concentration at which the maximal number of cross-links can be formed and that this optimal cross-linker concentration is for our model parameters at a half-equimolar to equimolar cross-linker-to-lysine concentration. This ratio is in excellent agreement with existing recommendations based on experimental observations (*see* for example ^17^) but could only now be derived from first principles.

In summary, our model based on chemical kinetics allows us to select tunable parameters and to find optimal conditions for cross-link formation. This way it enables us to operate prospectively under an experimental regime where cross-links can be maximized. It also helps to pave the way for the design of future experiments and cross-linkers that have the potential to generate a maximal number of protein-protein cross-links for the investigation of how proteins interact and communicate within the cell.

## Materials and Methods

### Data Availability

The code for the creation & evaluation of the kinetic model is available at:

https://github.com/stengel-laboratory/xlink-kme-sbml The simulation results are available at:

https://github.com/stengel-laboratory/xlink-kme-sbml-analysis

### Kinetic Model

The simplest kinetic model consists of a cross-linker that can either hydrolyze and/or react with two lysines. The products of the reaction are the mono- or bihydrolized cross-linker, a mono-link with lysine 1, a mono-link with lysine 2 or a cross-link between lysine 1 and 2 (**Fig 1**). This model may be expanded to any number of lysines with the 7 distinct species included (**Supp Table 2**). We obtain 10 free parameters, including 2 reaction rates, 6 diffusion rates and 2 concentrations (**Supp Table 3**).

The number of reactions that one must define increases exponentially with the number of lysines if there is no distance cut-off for the cross-link reaction. Therefore, if the structure for a given protein is known, distances higher than a defined maximum should be ignored. We developed a framework that generates all the species, and builds the corresponding kinetic equations, for any given protein complex. For our models we chose a cutoff at 30 Å for the solvent-accessible surface distance (SASD)^32^.

We use a transition-state model for the acylation reaction, where a diffusion-limited process brings the cross-linker in a reaction-competent state with a rate *k_on,i_*, and a rate *k_off,i_* to dissociate from it. The index *i* denotes a specific lysine. For simplicity, we assume the same bulk diffusion rates for all lysines; i.e. *k_on,i_*= *k_on_* and *k_off,i_* = *k_off_*. In the same fashion, we assume the same diffusion rates for the hydrolyzed and non-hydrolyzed cross-linker so that *k_on_^h^* = *k_on_*.

For our simulations, *k_on_* is approximated as the diffusion rate of a molecule the size of DSS in water at 20 °C^33^.

*k_h_* represents the hydrolysis rate of the cross-linker. It is a first order rate constant; other than *k_h_* itself, the hydrolysis rate depends on the cross-linker concentration. Experimentally, this will mostly depend on temperature and pH.

*k_lys,i_* is the intrinsic reactivity of each lysine. It will depend on factors like local chemical environment, pKa and accessibility. For our simulations, *k_lys,i_* is approximated for each lysine based on pKa and the solvent-accessible surface area which is calculated from the protein structure.

*k^XL^_on,i,j_* is the lysine-pair specific diffusion rate for a mono-link between these two lysines. It is the kinetic representation of the distance between lysines, which we call the *kinetic distance*. We derive it from the SASD distribution between lysine pairs calculated from the structure of a given protein, which is then scaled to a diffusion rate distribution.

In the following, the different aspects of the model are explored in detail.

## Hydrolysis

As soon as it is in solution, the cross-linker readily hydrolyzes on either end, leading to the same bihydrolyzed product, owing to the symmetry of the linker:

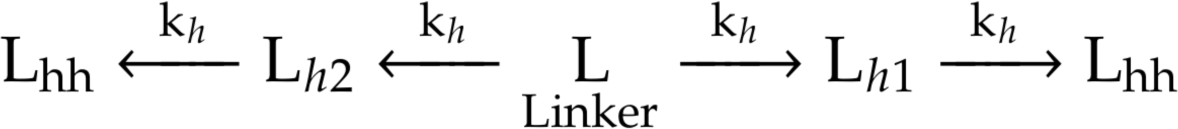

The differential equations describing this reaction, which do not include any reaction with a lysine yet, are as follows:

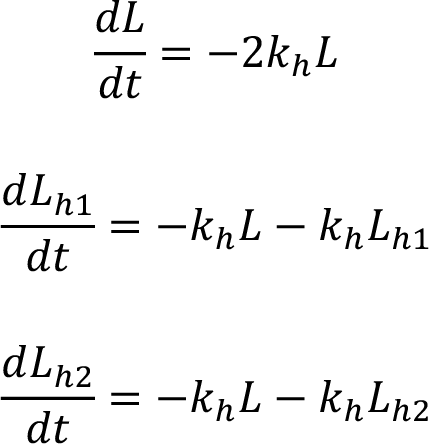

In practice, one cannot differentiate which end of the linker hydrolizes. Additionally, the concentration of either monohydrolyzed species should be equal at any given time, so that *L_h_* = *L_h1_* = *L_h2_*. Therefore,

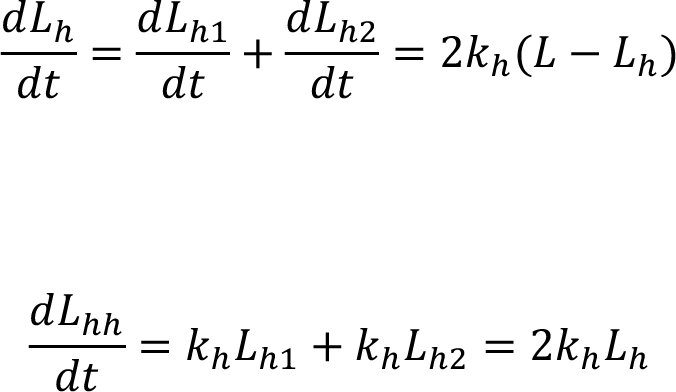

Note that all species symbols represent their time-dependent concentration. We find the following solutions to these equations:

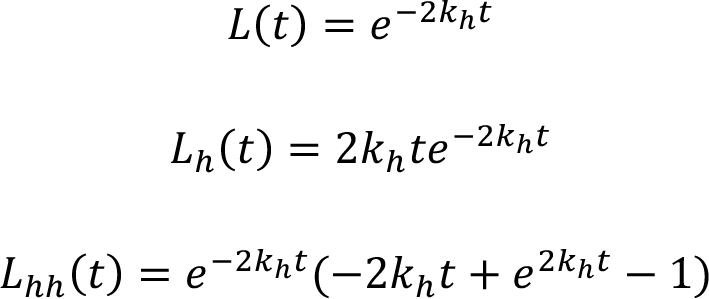

These equations describe the concentration of the different cross-linker species over time when considering hydrolysis as the only reaction. The cross-linker is depleted by hydrolysis while the mono-hydrolyzed cross-linker passes through a maximum before conversion to the bi-hydrolyzed cross-linker (**Supp Fig 7**).

## Diffusion and Acylation

In a kinetic model any diffusion process can be described as a reversible reaction. To encounter each other, the reactants must move through the solvent. This diffusion-limited event has its own activation barrier due to the viscous nature of the solvent and the entropic cost of bringing together the reactants. Once the reactants are in close proximity, they are still surrounded by the solvent that forms a *cage* around them^34^.

The reactants may either overcome the activation barrier to form a product or diffuse out of the solvent cage without reacting (**Supp Fig 8**).

The diffusion time needed to access the solvent cage is limited by *k_on_* while leaving the solvent cage is limited by *k_off_*.

The diffusion of a free cross-linker towards a lysine and the subsequent acylation is governed by the following equation:

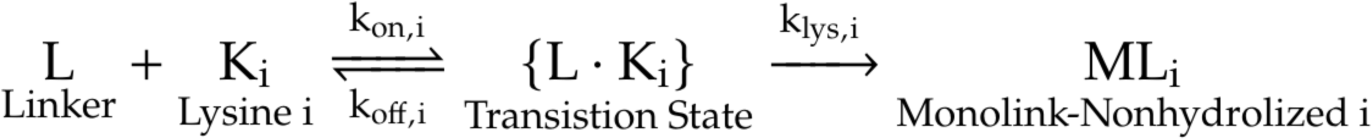

Likewise, the diffusion of a mono-hydrolized cross-linker and the following reaction is:

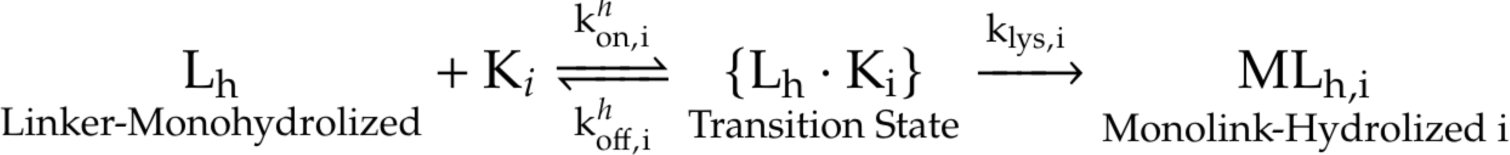

Here, we assume the same lysine-specific reaction rate constants *k_lys,i_* for both the non-hydrolized and the mono-hydrolized linker. We also assume that the diffusion of the non- and monohydrolized cross-linker feature the same rate constants for the forward diffusion, so that *k_on,i_*= *k^h^_on,i_*. *k_on,i_* describes the diffusion of a free cross-linker molecule in the bulk solution towards a specific lysine. The non-hydrolized cross-linker is bi-functional with two indistinguishable reactive groups. This is a simplification, as the local concentration of reactive groups for a non-hydrolized cross-linker molecule will be higher compared to that of a monohydrolized cross-linker. For the diffusion out of the solvent cage without a reaction, we assume that there is no discernible difference between the two rates, such that *k_off,i_* = *k^h^_off,i_*.

Since lysine side chains inside a protein are located within complex 3D structures, *k_on,i_* is lysine specific as it incorporates the accessibility of a lysine. Here, we assume that the accessibility is implicitly included in the lysine reactivity *k_lys,i_*. Therefore, *k_on,i_* will have the same value for all lysines *k_on_* = *k_on,i_*, and the same then also holds true for *k_off_*: *k_off_* = *k_off,i_*.

The non-hydrolized mono-link can further react to form a cross-link:

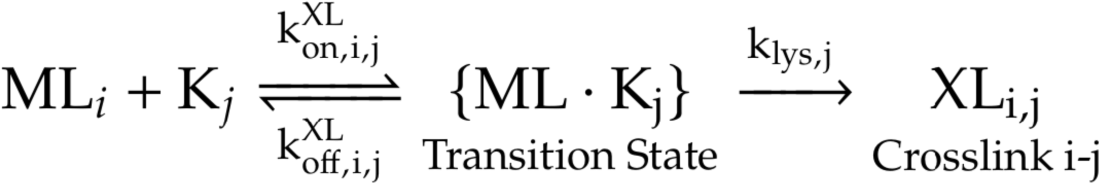

The diffusion processes of mono- and cross-link formation look similar but are governed by different diffusion rates, *k_on_* and *k^XL^_on,i,j_*. Since there is no a-priori knowledge concerning *k^XL^_off,i,j_* we assume a single value, so that *k^XL^_on,i,j_* = *k_off_*. *k^XL^_on,i,j_* corresponds to the diffusion of a non-hydrolized mono-link, covalently bound to a protein (**Supp Fig 9**). Thus, we refer to *k^XL^_on,i,j_*as the *kinetic distance of the two lysines i and j*.

Importantly, *k^XL^_on,i,j_* will depend on the solvent-accessible surface distance (SASD) between the mono-link and the lysine. It cannot be simplified to a single parameter as done with *k_on_* as it will depend on the distance of each lysine pair. Ultimately, *k^XL^_on,i,j_*connects the structure of a protein to the cross-link concentration by combining both accessibility and distance between a lysine pair into a rate constant.

*k_lys,i_* then represents the individual reactivity of a given lysine which will depend on factors like the chemical environment and accessibility and is identical to the parameter used for the mono-link formation. Lysine reactivity can span several orders of magnitude within the same protein^35–37^. Note that all cited studies do not deconvolute diffusion and reactivity, they rather report an effective rate constant, as discussed in the next section.

### Simplified Kinetic Model

The main difference between the full (**Supp Fig 1**) and simplified kinetic model is the treatment of diffusion. In the full model diffusion is explicitly considered, whereas in the simplified model it is integrated in the rate constants of the chemical reactions. Furthermore, the simplified model includes the approximations that are introduced in the section *Diffusion & Acylation* for the full model.

In actual experimental measurements, diffusion is usually treated implicitly and included in an effective reaction rate constant. In our case, we introduce the parameters *k_eff,i_* and *k^XL^_eff,i,j_* as effective rate constants for the mono- and cross-link reaction.

First, consider a general diffusion reaction:

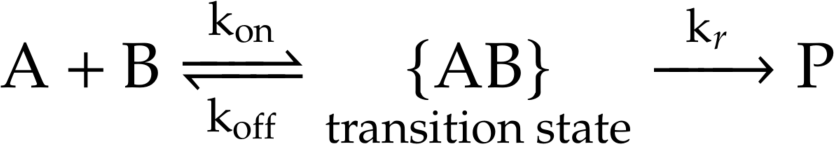

The concentration change over time of the product P is

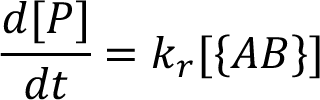

For the transition state the time-dependent concentration change is

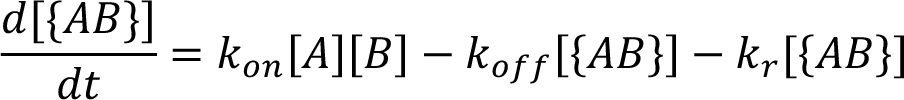

When assuming steady-state, i.e. 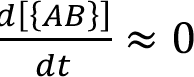, we can simplify as follows:

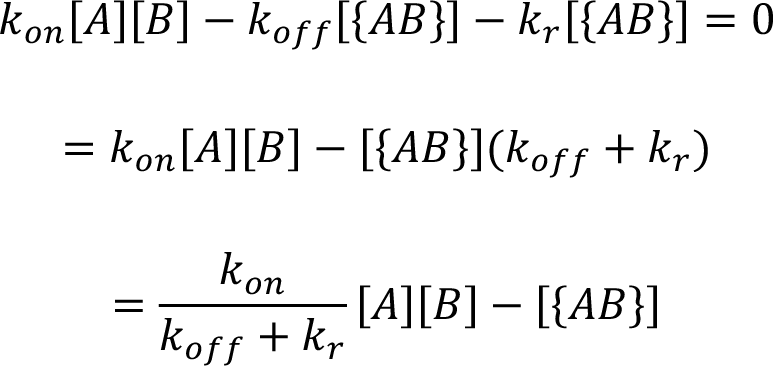

Therefore,

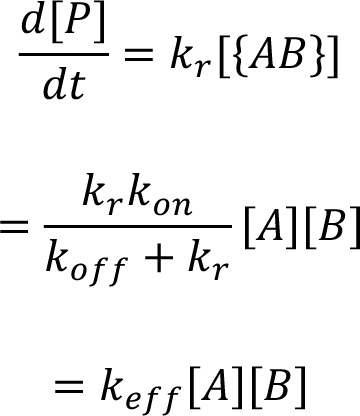

The effective reaction rate is thus described by

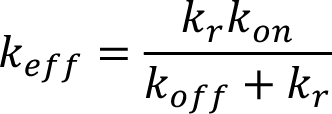

Coming back to our general diffusion reaction this leads to

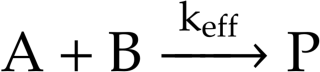

We further explore the case of an activation-controlled reaction: here, the reactants require energy from the surrounding liquid to overcome the activation barrier. Therefore, only very few encounters of the reactants lead to a product (𝑘_*r*_ ≪ *k_off_*). This is a reasonable assumption for our reaction as opposed to a diffusion-controlled reaction where every encounter between species leads to a product.

Therefore, in the case of activation-control, the effective rate constant can be simplified to

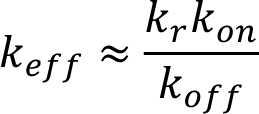

In this case both the activation barrier of moving the reactants through the solvent as well as the activation barrier of the reaction itself determine the effectively measured reaction rate.

In summary, we arrive at a simplified kinetic model with the following reactions for two lysines:

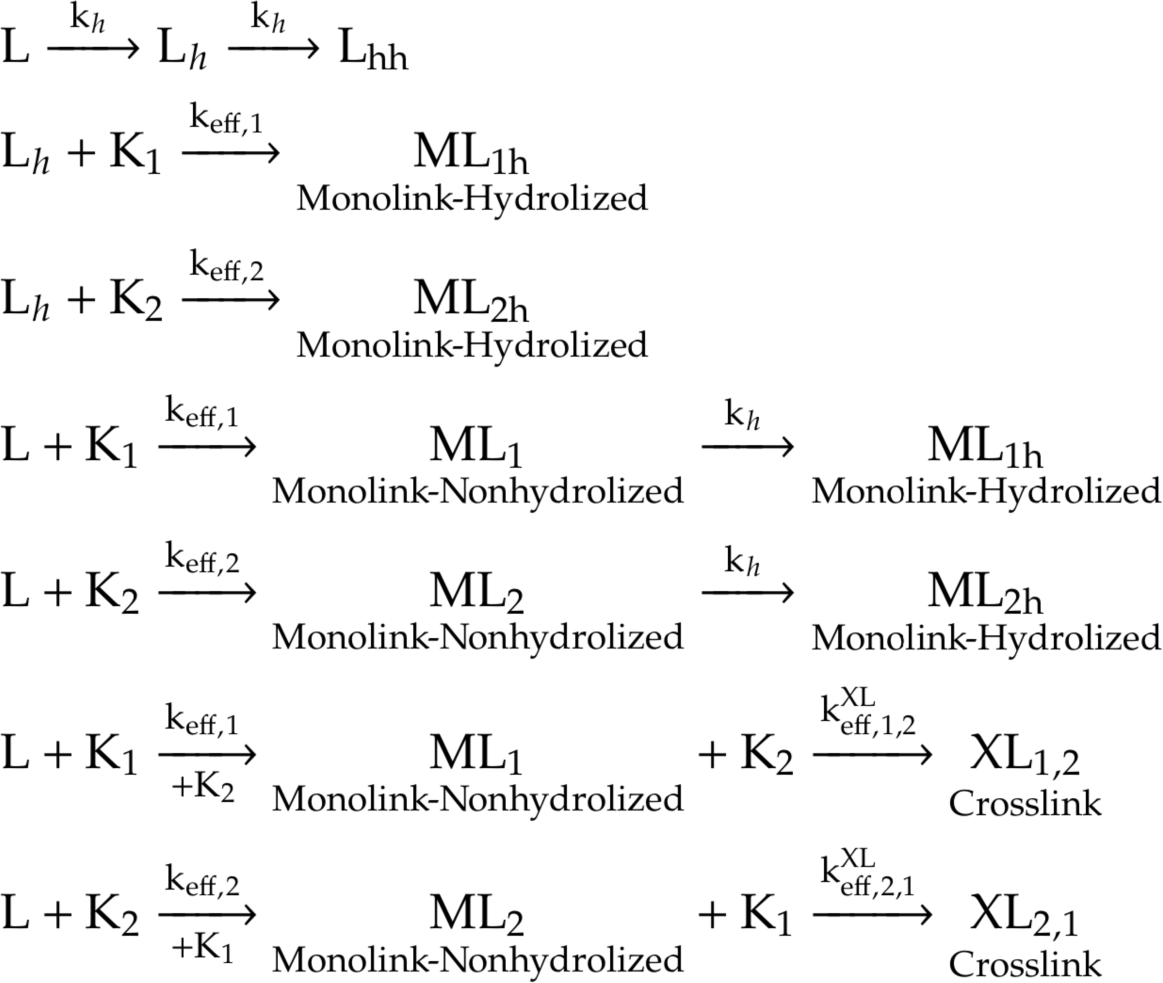

With

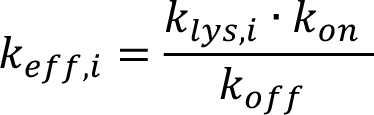

and

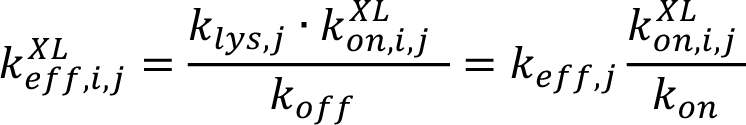

Since we directly estimate *k_eff,i_* from a protein structure (*see* section **Model Builder**), we can eliminate *k_off_* as input parameter and aggregate it with others (**Supp Table 4**). The resulting *effective kinetic distance k^XL^_eff,i,j_* is a composite rate constant containing the product of the kinetic distance and the effective lysine reactivity divided by the cross-linker diffusion rate.

### Kinetic Simulation

Every kinetic simulation consists of three main steps:

1. kinetic model definition and export
2. numerical simulation
3. analysis and visualization

For every system (i.e. each protein or protein complex), a new model needs to be created. All models below are based on the simplified kinetic model introduced above and then adapted to a specific system. The models are created by a framework facilitating the Systems Biology Markup Language (SBML)^38^, a language designed for creation and exchange of biochemical models. The numerical simulation is done by Tellurium^39,40^ which can natively import the SBML models. Finally, simulation results are analyzed using a mixture of a Python backend and an iPython notebook frontends.

The values for the parameters used in our simulation orient themselves on known literature values^35,41,42^ and are adjusted to obtain reasonable amounts of significant cross- and mono-links at a selected significance cut-off at 0.05 g/L (**Supp Table 1**). Simulation results concerning relative changes are set to use an absolute concentration change of 0.05 g/L and a log2ratio of at least 0.5 for significant changes.

We ran simulations of a kinetic model for cross-linking based on information gathered from protein structures. The simulations are extended until complete hydrolysis of the sample, which is a sign for the completion of the reactions.

### Model Builder

The kinetic model of a protein complex is facilitated by information provided by its structure. The model builder includes a suite of different python scripts. Two scripts will estimate local lysine pKa (via pKAI^43^) and the solvent-accessible surface area (SASA, via Biopython^44^). Another script will prepare the output of solvent-accessible surface distance (SASD) between lysine pairs obtained via jwalk^45^. Taken together, the output prepared by these three scripts serves as input for the model builder, containing estimated lysine reactivities as well as kinetic distances (**Supp Fig 10**).

The model builder is the user front-end for the kinetic framework. It will create a SBML cross-linking model using the provided input. It will normalize and scale the physical input parameters like pKa, SASA and SASD to plausible ranges for the corresponding kinetic parameters of the model (**Supp Fig 11** and **Supp Fig 12**). This is also the place, where other parameters like hydrolysis and cross-linker diffusion rate are defined. Besides the SBML model itself, the model builder also outputs all model parameters as a csv file. These model parameters may be modified and then re-used as an input for the model builder. The SBML model is subsequently used as input for the simulation framework.

### Cross-link Simulation Framework

With the SBML model in place, any simulation may now directly be run without any further modification in Tellurium. The output of the simulation is then the time-dependent concentration of all species (e.g., the concentration of a cross-link or a mono-link at a specific site). For most simulations it is insightful to vary model parameters across a simulation (e.g. modifying hydrolysis, lysine reactivity or cross-link kinetic distance). To do so, we created a flexible framework which allows for systematic modifications applied to a model before simulating it with Tellurium. It is of note though that the simulation time scales exponentially with the number of parameters. The framework also defines and writes the output file of a simulation in pandas^46^ data structures, with a multitude of detailed information to facilitate post-simulation analysis.

### Simulation Evaluation & Visualization

The results obtained by simulation are evaluated and visualized using the Python library altair^47^, using a combination of Python scripts and Python notebooks (the user front-end). There are a multitude of tools to exhaustively analyze a simulation. Convergence can be assessed as well as the distribution of the different species. It is possible to visualize the concentration of every single species over time or for the final frame of the simulation.

Also, analysis of aggregates like the mean, median or overall quantity of a species is readily available. Furthermore, there are options to visualize the influence of any kinetic parameter on the concentration or the influence of changing two parameters at the same time. Finally, the quantity of mono- and cross-links can be mapped on the protein structure and displayed by ChimeraX^48^. All the plots in this work were created using the scripts described above.

## Author Contributions

K.M.K, R.P. and F.S conceived the study and experimental approach. K.M.K & R.P. designed the kinetic model, performed kinetic analyses and simulations. K.M.K wrote and implemented all code with help from R.P. K.M.K, R.P. and F.S. analyzed the data. K.M.K, R.P. and F.S wrote the paper.

## Classification

Biological Sciences (Biophysics and Computational Biology), major classification; Physical Sciences (Chemistry), minor classification.

## Acknowledgments

This work was supported by the German Research Foundation, project numbers 496470458, 684699 and TRR353, and the International Emerging Action (IEA) of CNRS 2019-2020.

