## Supplementary Information for "Kinetic principles of chemical cross-link formation for protein-protein interactions"

**This PDF file includes:**

Figures 1 to 12

Tables 1 to 4

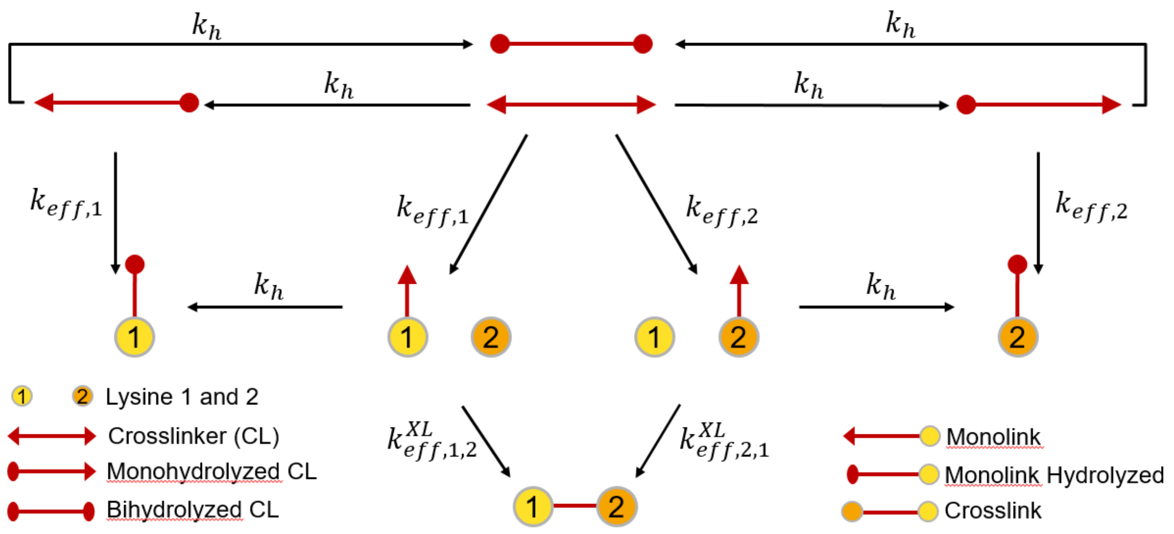


**Supp Fig 1. Simplified kinetic model.** A two-lysine model detailing the reactions of a cross-linker taking hydrolysis, cross-link and mono-link formation into account. Diffusion is treated implicitly.

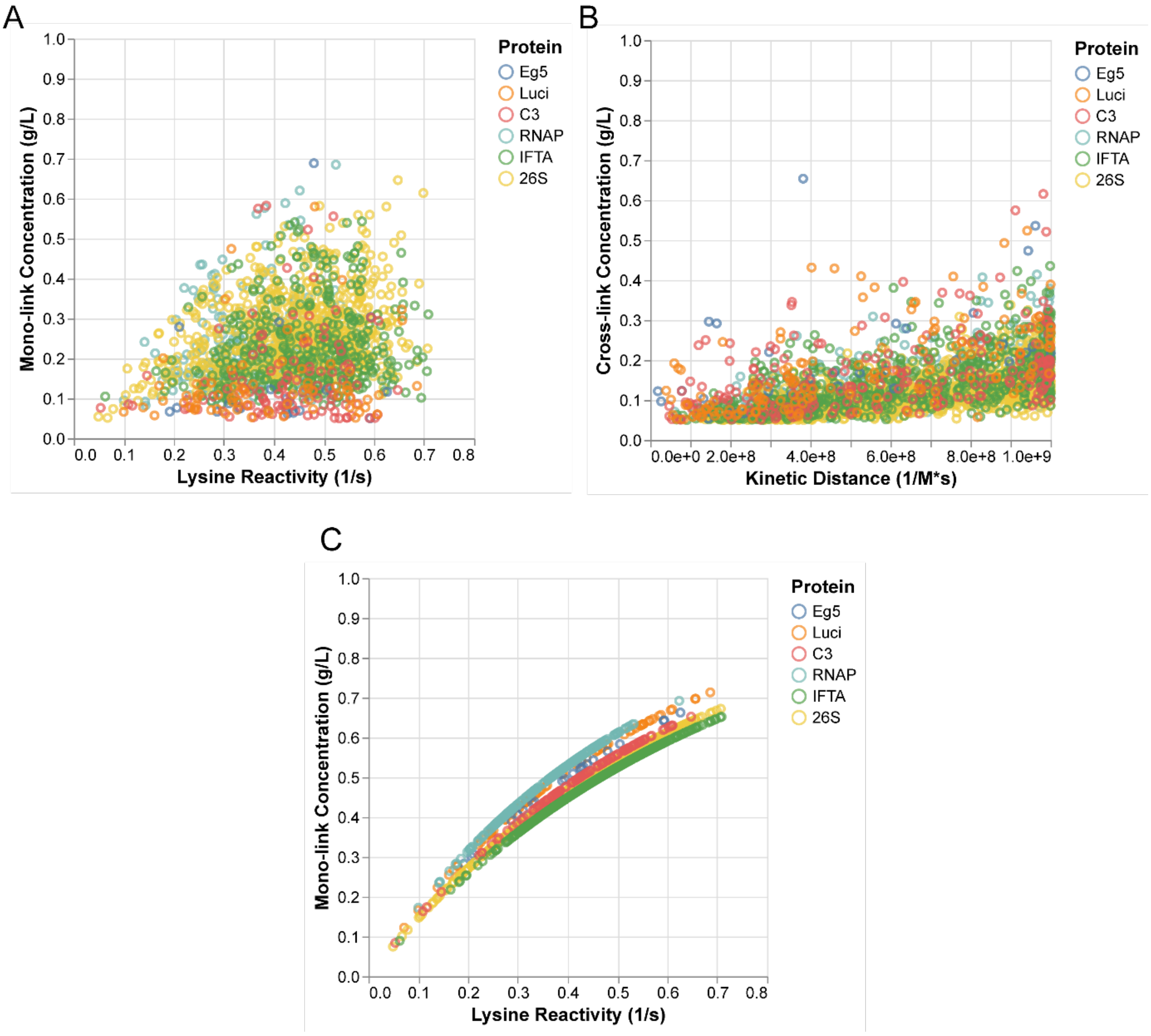


**Supp Fig 2. Correlation between rate constants and mono-/cross-links. A** Plotted is the final concentration of mono-links versus the lysine reactivity. **B** Plotted is the final concentration of cross-links versus the kinetic distance (*see* Materials and Methods for details). **C** Plotted is the final concentration of mono-links versus the lysine reactivity for the “mono-link-only” model.

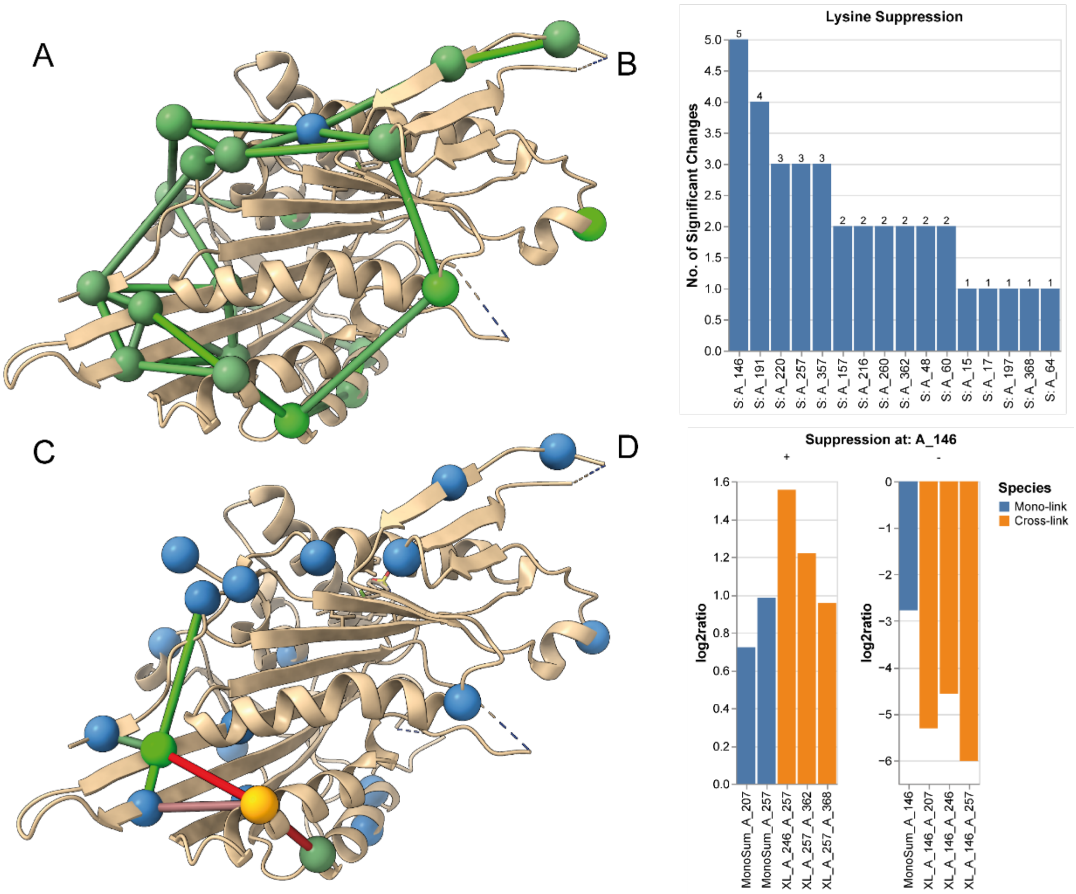


**Supp Fig 3. Effect of lysine reactivity suppression on cross-link formation**. **A** Structure of Eg5 showing the formed mono- (green beads) and cross-links (green bonds) as obtained from the kinetic simulation. The color intensity is proportional to the final concentration of the formed mono- or cross-link. Non-significant mono-links are shown in blue. **B** The reactivity of each lysine in the Eg5 protein is successively set to zero, such that it can form no longer mono- or cross-links, before the corresponding model is simulated. Significantly (see methods) up or down regulated mono- and cross-links are identified by comparison to the reference dataset (i.e., the simulation in panel **A**). For example, there are 5 significant changes resulting from the suppression of lysine at position 146 in chain A (first blue bar). Note that "trivial" down-regulations, i.e. links that are directly associated with the given lysine position are not shown, e.g. when suppressing lysine 146, the mono-link at position at lysine 146 and the cross-links directly connected with this lysine are not considered. **C** Structure of Eg5 showing the significant up- and down- regulated mono- and cross-links after suppression of lysine at position 146 in chain A. The suppressed lysine is shown as a yellow bead and non-regulated lysines are shown as blue beads. Significant up regulations are shown in green and down regulations in red and colored according to their intensity. Note that all down regulations have a direct connection to the suppressed lysine and therefore count as "trivial" changes (see panel **B**). **D** Significantly changed mono- (orange) and cross-links (blue) resulting from the suppression of lysine A146. Note that all down regulations have a direct connection to the suppressed lysine and therefore count as "trivial" changes (see panel **B**).

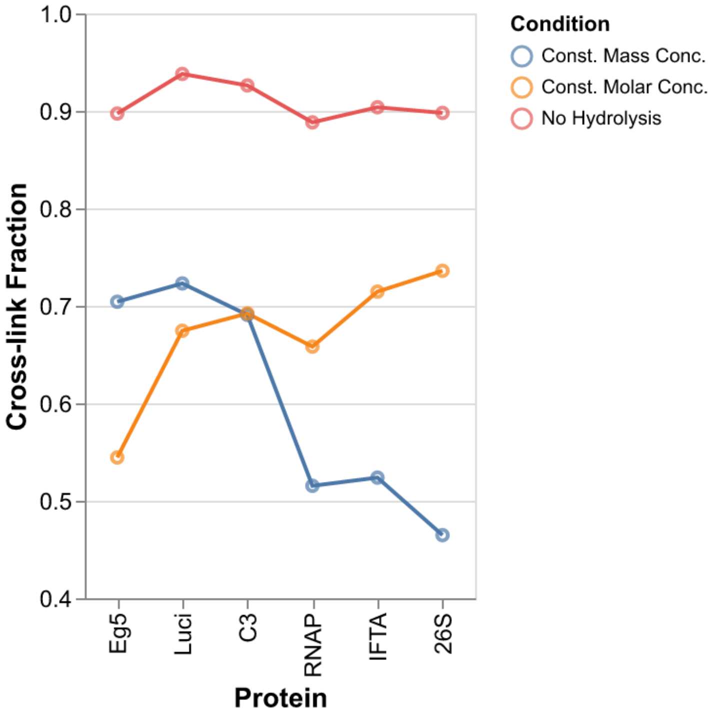


**Supp Fig 4. Cross-link fraction depends on protein concentration.** Plotted is the cross-link fraction of the six model proteins. The cross-link fraction is calculated by dividing the sum of all cross-links by the sum of all mono- and cross-links (y-axis). Proteins are ordered by size, from smallest (Eg5) to largest (26S) (x-axis). Three conditions are shown: constant mass concentration in blue (protein concentration at 1 g/L), constant molar concentration in orange (protein concentration at $5.5 \cdot 10^{-6} \frac{mole}{liter}$) and a condition without hydrolysis and constant mass concentration in red (protein concentration at 1 g/L). Other parameters use the default values (**Supp Table 1**).

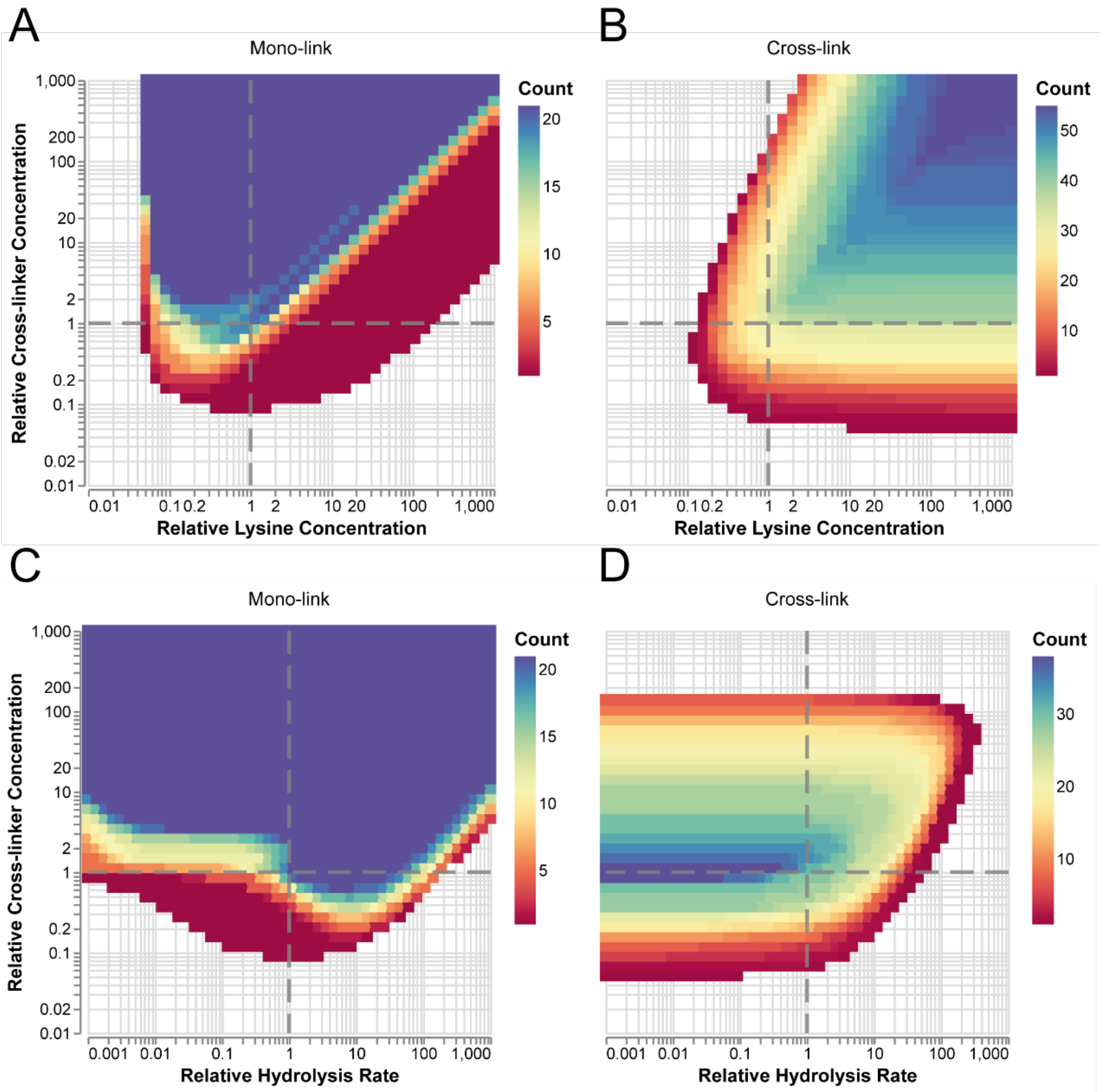


**Supp Fig 5. Mono- and cross-link formation as a multi-parameter function (Eg5)**. First row: relative cross-linker concentration plotted versus the relative mean lysine concentration. Second row: relative cross-linker concentration plotted versus the relative hydrolysis rate. The color corresponds to the number of formed mono-links (**A**, **C**) and cross-links (**B**, **D**) identified for the given parameter combination using an intensity cut-off of 0.05 g/L. The reference value of each parameter is indicated by the dashed lines (**Supp Table 1**).

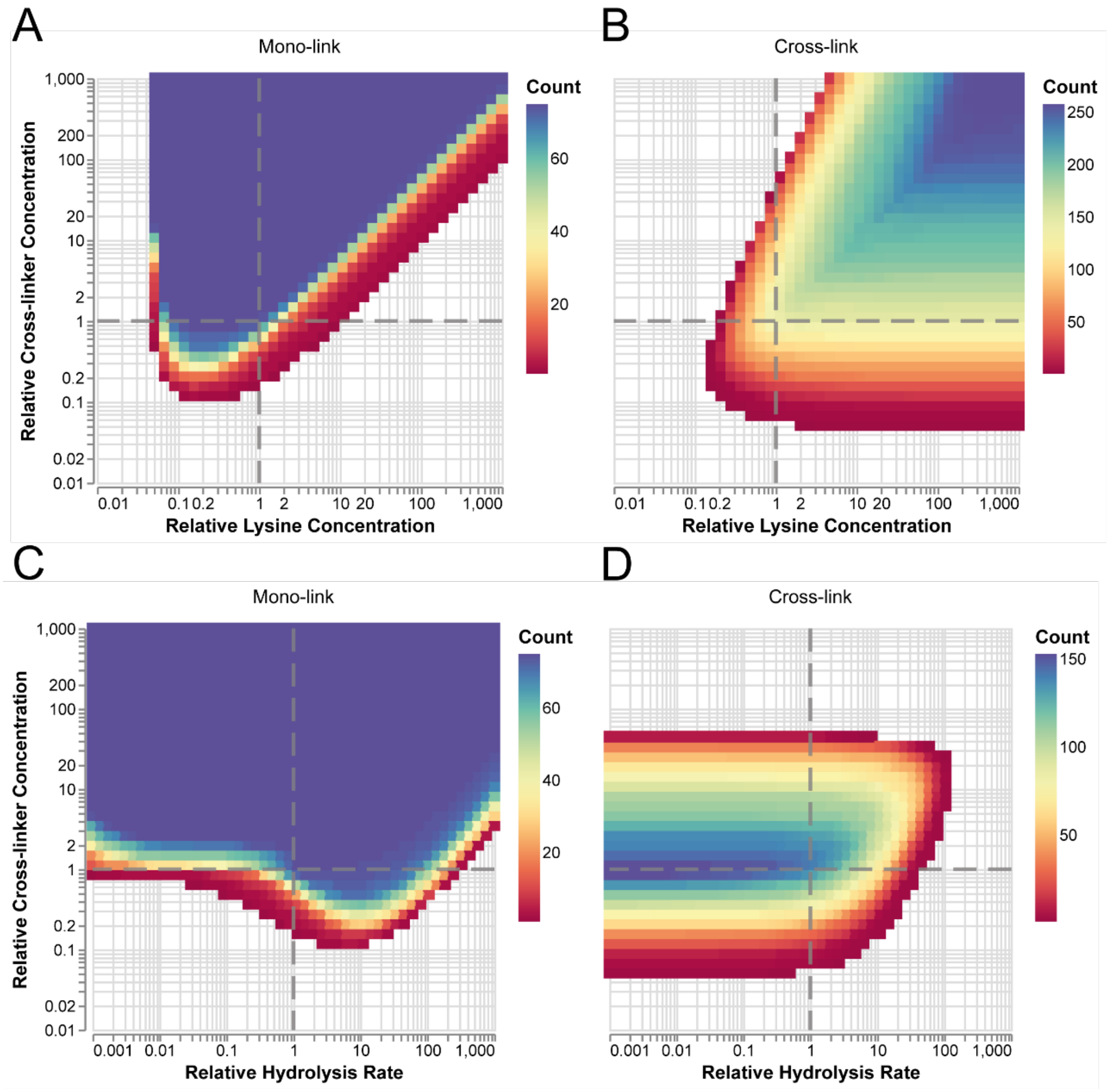


**Supp Fig 6. Mono and cross-link formation as a multi-parameter function (Luciferase)**. First row: relative cross-linker concentration plotted versus the relative mean lysine concentration. Second row: relative cross-linker concentration plotted versus the relative hydrolysis rate. The color corresponds to the number of formed mono-links (**A**, **C**) and cross-links (**B**, **D**) identified for the given parameter combination using an intensity cut-off of 0.05 g/L. The reference value of each parameter is indicated by the dashed lines (**Supp Table 1**).

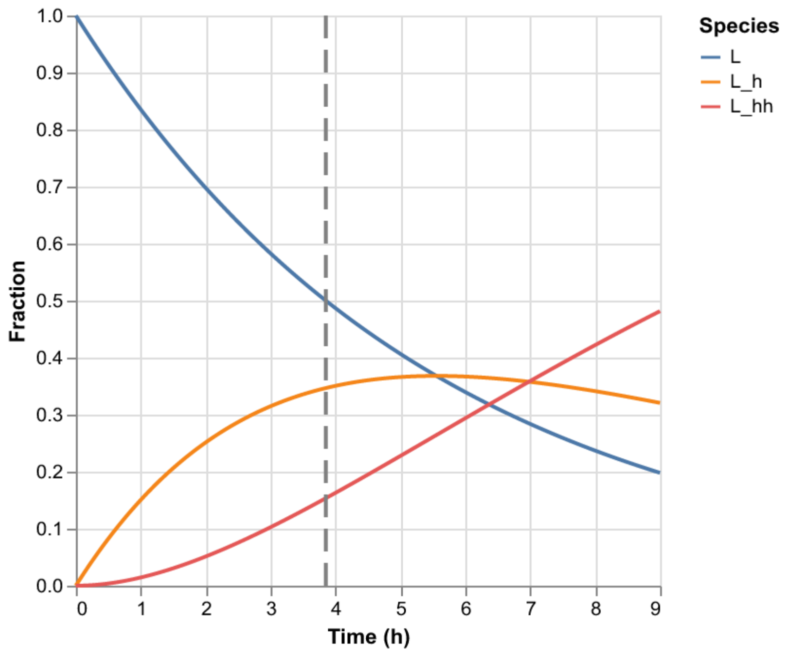


**Supp Fig 7. Cross-linker hydrolysis.** Visualizing the analytical solution to the hydrolysis part of the kinetic model. The half-life of the cross-linker is marked with a dashed gray line at ~4 h. The hydrolysis rate presented in **Supp Table 1** is used. The abbreviations shown in the legend are explained in **Supp Table 2**.

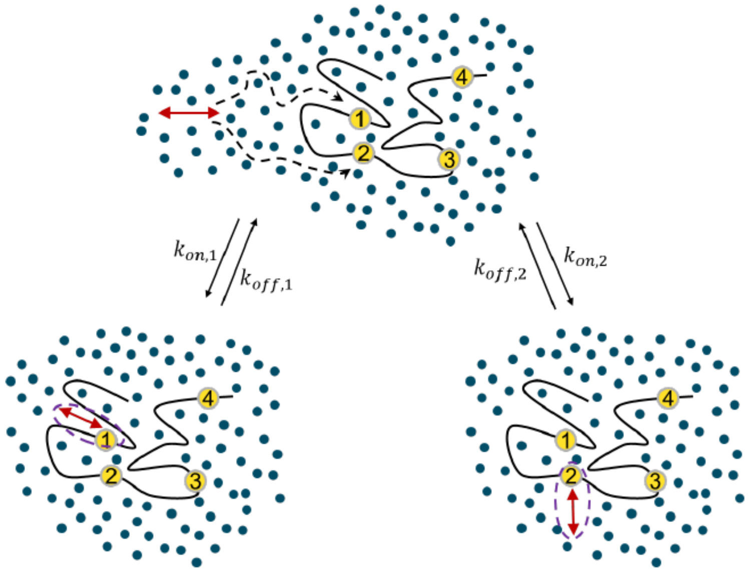


**Supp Fig 8**. **Cross-linker diffusion.** The cross-linker diffuses either towards lysine 1 or lysine 2. It collides with the solvent (blue dots) which forms a solvent cage around the transition state (purple). Lysine 1 and 2 have a different accessibility, reflected by their reactivities.

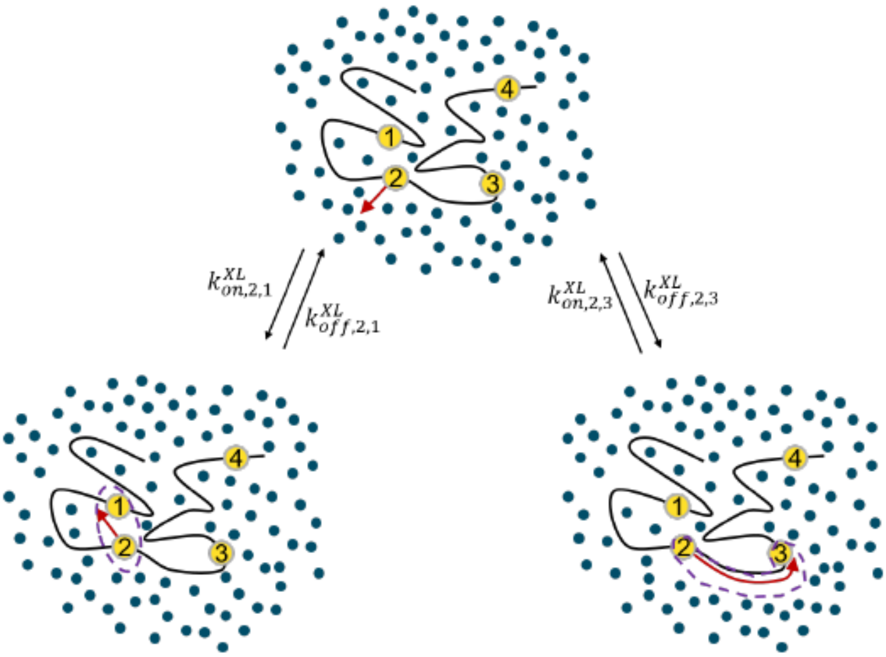


**Supp Fig 9. Mono-link diffusion to form a cross-link.** The mono-link either diffuses towards lysine 1 or lysine 3 (lysine 4 is not within reach of the cross-linker). The SASD to lysine 1 is much shorter than that to lysine 3 which is reflected in the respective rate constants (kinetic distance).

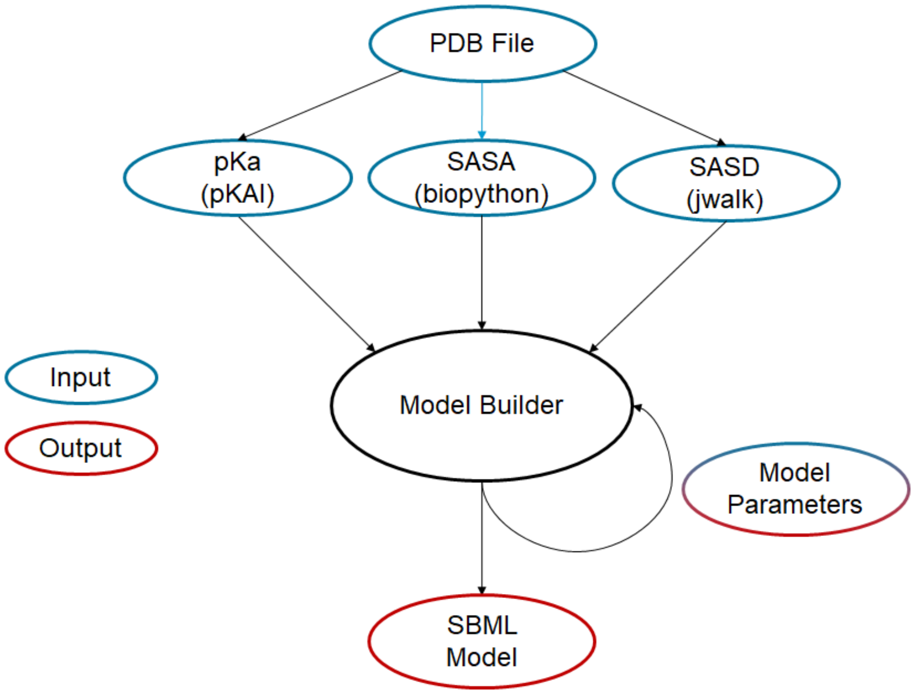


**Supp Fig 10. Kinetic Framework Overview.** A single PDB file is the only input for the protocol. Kinetic parameters are inferred from the PDB file: lysine reactivities from pKa values and solvent-accessible surface area (SASA), while the kinetic distance is computed from the solvent-accessible surface distance (SASD). A SBML model is generated as output as well as a dataframe containing the model parameters ("Model Parameters"). Parameters in the dataframe may be modified and then used as input again. The output is a xml file, which may be read by any software compatible with the SBML format.

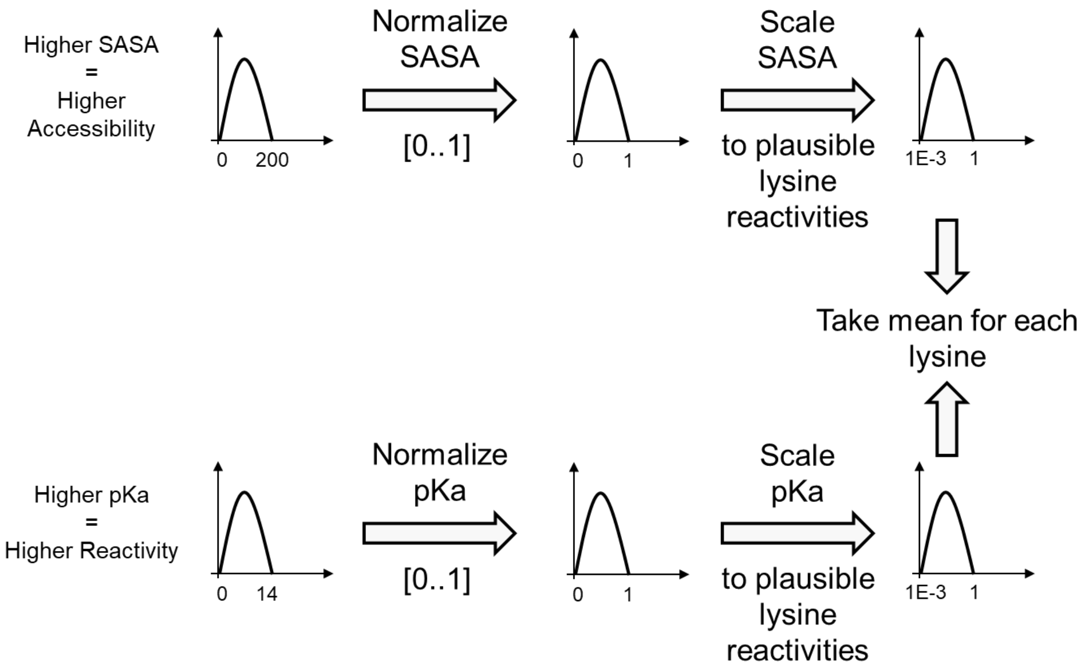


**Supp Fig 11**. **Calculation of lysine reactivity from the structure.** SASA and pKa are extracted from the structure of the system, and normalized in a range of [0,1]. They are then scaled to plausible ranges for lysine reactivities and the mean of the two values is taken as lysine reactivity.

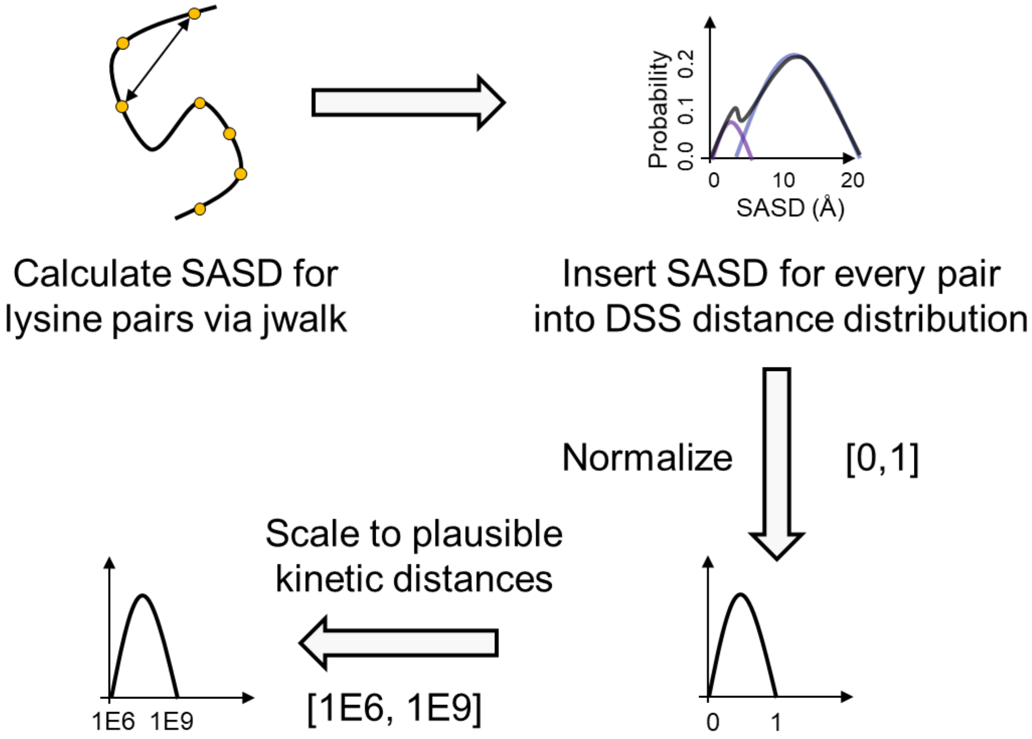


**Supp Fig 12**. **Calculation of the kinetic distance from the crystal structure.** The SASD for each lysine pair is extracted from crystal structure and fed into the expected distance distribution of DSS cross-linked lysine residues[1]. Next, the distribution is normalized to a range of [0,1] and finally scaled to plausible ranges for kinetic distances which we assume to be around the diffusion rate of DSS in water.

**Supp Table 1.** The default parameters used for the kinetic simulations. Note that $k_{eff,i,j}^{XL}$ is oriented at the diffusion rate of DSS in water (*see* **Materials & Methods**).

| Parameter | Value | Unit | Reference |
|---|---|---|---|
| $simulation\ time$ | $1.0 \cdot 10^{7}$ | $s$ | - |
| $LYS$ | $1$ | $\frac{g}{L}$ | - |
| $XL$ | $0.5 \cdot number\_lysines$ | $\frac{g}{L}$ | - |
| $k_h$ | $2.5 \cdot 10^{-5}$ | $s^{-1}$ | 2,3 |
| $k_{on}$ | $1.0 \cdot 10^{7}$ | $M^{-1}s^{-1}$ | 4 |
| $k_{eff,i}$ | $1.0 \cdot 10^{-3}\ to\ 1.0$ | $M^{-1}s^{-1}$ | 2,3,5 |
| $k_{eff,i,j}^{XL}$ | $1.0 \cdot 10^{6}\ to\ 1.0 \cdot 10^{9}$ | $M^{-1}s^{-1}$ | - |
| $k_{eff,i}$ (Three Lysine Sim.) | $0.5$ | $M^{-1}s^{-1}$ | 2,3,5 |
| $k_{eff,i,j}^{XL}$(Three Lysine Sim.) | $9.0 \cdot 10^{7}$ | $M^{-1}s^{-1}$ | - |

**Supp Table 2**. Species and their abbreviations used in the kinetic model.

| Species | Abbreviation |
|---|---|
| Lysine | $K$ |
| Crosslinker | $L$ (Linker) |
| Crosslinker Monohydrolyzed | $L_h$ |
| Crosslinker Bihydrolyzed | $L_{hh}$ |
| Monolink | $ML$ |
| Monolink Hydrolyzed | $ML_h$ |
| Crosslink | $XL$ |

**Supp Table 3.** Parameters used in the kinetic model and their physical meaning.

| Parameter | Physical Meaning |
|---|---|
| $k_h$ | Hydrolysis |
| $k_{lys,i}$ | $K$ reactivity |
| $k_{on,i}$ | $L$ diffusion |
| $k_{on,i}^h$ | $L_h$ diffusion |
| $k_{off,i}$ | $L$ diffusion off ML transition state |
| $k_{off,i}^h$ | $L_h$diffusion off ML transition state |
| $\mathrm{k_{on,i,j}^{XL}}$ | $ML$ diffusion (kinetic distance) |
| $\mathrm{k_{off,i,j}^{XL}}$ | $ML$ diffusion off XL transition state |
| $LYS$ | Lysine concentration |
| $XL$ | Crosslinker concentration |

**Supp Table 4**. Parameters of the simplified kinetic model. The column to the right denotes which parameters from the full model are combined into the composite parameters of the simplified model.

| Parameter | Physical Meaning | Included Parameters from Full Model |
|---|---|---|
| $k_h$ | Hydrolysis | $k_h$ |
| $k_{eff,i}$ | Effective $K$ reactivity | $k_{lys,i}$, $k_{on,i}$, $k_{on,i}^{h}$, $k_{off,i}$, $k_{off,i}^{h}$ |
| $\mathrm{k}_{eff,\mathrm{i,j}}^{\mathrm{XL}}$ | Effective XL reactivity | *See* $k_{eff,i}$ plus $\mathrm{k}_{\mathrm{on,i,j}}^{\mathrm{XL}}$, $\mathrm{k}_{\mathrm{off,i,j}}^{\mathrm{XL}}$ |
